# Characterization of effects of a neurotropic murine coronavirus infection on Alzheimer’s disease neuropathology of 5xFAD mice

**DOI:** 10.64898/2026.02.23.707587

**Authors:** Dominic I. Javonillo, Susana Furman, Lucas Le, Kellie Fernandez, Jazmyn Mulford, Vidushi Singla, Roshni Jha, Kate Inman Tsourmas, Nellie E. Kwang, Kim N. Green, Thomas E. Lane

## Abstract

**Background:** Recent studies revealed key immunological mechanisms within the central nervous system (CNS) that contribute to Alzheimer’s disease pathology. Additionally, analyses of human AD datasets have also associated viral encephalitis exposure (i.e., viral-induced neuroinflammation) with the development of AD and dementia, highlighting the need to better understand how viral encephalitis and neuroimmune mechanisms within the brain may impact AD pathologies such as Aβ plaque deposition. Intracranial infection of susceptible mice with the neurotropic JHM strain of murine coronavirus (JHMV) results in acute encephalomyelitis characterized by viral infection of glia and a robust inflammatory response comprised of monocytes/macrophages and T cells that aid in controlling viral replication.

**Methods:** To determine how coronavirus-induced encephalitis may impact established Aβ plaque deposition, we intracranially inoculated JHMV into aged 5xFAD model of amyloidosis. We utilize immunohistochemical and biochemical analysis to assess the impact on existing Aβ pathology. We also utilize spatial transcriptomic imaging to explore how viral encephalitis affects cellular responses to plaque pathology with single-cell resolution.

**Results:** In aged 5xFAD mice, JHMV-induced encephalitis at 12 days p.i. resulted in minimal changes to overall Aβ protein within the brain. However, viral encephalitis induces CD4^+^ and CD8^+^ T cell infiltration and more *Lgals3/*MAC2-expressing macrophages surrounding dense-core Aβ plaques, which appear more compacted in JHMV-infected 5xFAD brains compared to uninfected 5xFAD controls. We compared gene expression within JHMV-infected 5xFAD mice and uninfected controls to identify distinct cellular responses to Aβ plaques that differed. Utilizing differential gene expression and pathway analysis, we found that viral encephalitis increased the proportion of myeloid cells in the 5xFAD brain, which also showed down-regulated disease-associated (DAM) pathways involving Aβ clearance, response to lipids, and macrophage activation within the post-encephalitis 5xFAD brains.

**Conclusions:** Together, these findings suggest an attenuated myeloid cell response to Aβ plaque burden in 5xFAD mice following acute viral encephalitis. Future experiments aim to further dissect inflammatory mechanisms between infiltrating myeloid cells, T cells, and the progression of Aβ and tau pathology. Data derived from these experiments will further elucidate the viral-induced neuroimmune mechanisms that affect AD pathology and offer an opportunity to determine how these neuropathologic changes, such as subsequent neuronal damage, occur.

## Background

Alzheimer’s Disease (AD) is the most common cause of dementia impacting almost 7 million people in the United States with cases projected to increase over 13 million by 2050^1,2^. As a progressive neurodegenerative disease, AD is defined by its substantial synaptic dysfunction and neuronal loss leading to brain atrophy, cognitive decline, and severe memory impairment^3,4^. Two characteristic hallmarks of AD pathology are the widespread deposition of neuritic beta-amyloid (Aβ) plaques and aggregation of neurofibrillary tau tangles (NFTs), which are proposed to occur consecutively or work synergistically towards neuronal damage and loss associated with AD^5,6^. As neurodegeneration progresses, cognitive impairments begin to manifest through subtle problems with memory, language, and thinking before escalating to significant impairments to behavior, motor, and executive functions^1^.

Over the past decade, genome-wide association studies (GWAS) have been a critical tool to determine polymorphic variants within genes that increase genetic risk and identify physiological processes contributing to AD pathogenesis in LOAD^7^. Numerous large GWAS have repeatedly identified AD risk genes associated with immunity and specifically expressed in myeloid cells, implicating neuroimmune processes contributing to altered neuroinflammation as a major factor of AD etiology. Myeloid cells within the CNS are primarily comprised of two distinct populations: CNS-resident microglia and monocyte-derived macrophages. Many studies have demonstrated a close association between Aβ plaques and microglia, which are considered the principal immune cell within the CNS. Once activated, microglia undergo a morphological and functional change into rounder, amoeboid shapes with short, thick processes that extend out to enable migration towards the inflammatory stimulus, cytokine secretion, and phagocytosis^8,9^.

Microbial infection has been broadly explored as a mechanism through which both genetic and environmental contributions interact together to influence risk of developing AD and impact ongoing AD neuropathology. Pathogenic microbes that invade the CNS can cause significant neuroinflammation through CNS-resident immune cells and infiltration of peripheral immune cells such as monocyte/macrophages and T cells. To illustrate the association of viral infection with several neurodegenerative diseases, Levine et al.^10^ mined medical record data from both the Finnish biobank (FinnGen) and UK Biobank. Their findings demonstrated several viral pathogens tightly associated with different neurological diseases, while the strongest association was between AD and viral encephalitis^10^. Viral encephalitis is defined as viral-mediated inflammation of the brain parenchyma. While the BBB prevents most viral pathogens from invading from the periphery, neurotropic viruses can circumvent these barriers to infect and replicate within the CNS^11^. The resulting inflammation via indirect neuroimmune mechanisms may induce acute and chronic neurologic damage with potential consequences in the onset or severity of neurodegenerative disease^12–14^. Therefore, it is unsurprising that majority of identified viral pathogens associated with neurodegenerative disease and dementia are neurotropic (i.e., herpes simplex virus (HSV), varicella zoster virus (VZV), Epstein-Barr Virus, and flaviviruses) and/or systemic infections leading to viral encephalitis (i.e., human immunodeficiency virus (HIV) and severe acute respiratory syndrome-Coronavirus 2 (SARS-CoV-2)^15–17^. Given the diversity of viral pathogens that can replicate within the CNS or instigate encephalitis, there is a varied inflammatory response that may contribute to the onset or severity of AD pathology. While the field’s current understanding of the contributions of these viruses and their resulting inflammation on AD remains nuanced and complex, ongoing investigations are precisely identifying direct and indirect viral mechanisms leading to CNS damage and associated cognitive impairments like dementia.

Our lab has employed experimental infection of mice with the neurotropic JHM strain of murine hepatitis virus (JHMV, a member of the *Betacoronaviridae* genus) to model molecular and cellular mechanisms of host defense and disease in response to viral encephalitis within the CNS^18,19^. Intracranial inoculation of susceptible mice with JHMV results in an acute encephalomyelitis characterized by wide-spread replication of virus in glia with relative sparing of neurons. Innate immune responses characterized by the expression of type I interferon are important in limiting the spread of virus early following infection. Ultimately, virus-specific CD4^+^ and CD8^+^ T cells are attracted into the CNS in response to T cell chemoattractant chemokines and control viral replication via expression of anti-viral cytokines including IFN-γ and cytolytic activity^20,21^. However, sterile immunity is not achieved and virus will persist within white matter tracts, where immune-mediated demyelination will occur due to chronic infiltration of macrophages and T cells^22,23^. Based upon evidence that neuroinflammation impacts AD-associated neuropathology and that viral encephalitis is now recognized as a major risk factor associated with AD, the overall objective of this investigation is to determine the molecular and cellular mechanisms by which viral-induced inflammation impacts AD pathology and neurodegeneration after the onset of plaque deposition.

## Methods

### Mice

Male and female C57BL/6 WT and 5xFAD at 6- and 10-months of age were used for these studies. Tg (APPSwFlLon, PSEN1*M146L*L286V) 6799Vas/ Mmjax RRID:MMRRC034848-JAX, was obtained from the Mutant Mouse Resource and Research Center (MMRRC) at The Jackson Laboratory, an NIH-funded strain repository, and was donated to the MMRRC by Robert Vassar, Ph.D., Northwestern University and have been previously described and characterized in detail^24,25^. In brief, the 5xFAD mouse expresses five familiar AD genes (APP Swedish, Florida, and London; PSEN1 M146L+L286V) and recapitulate progressive amyloid pathology as early as 3-4 months of age, leading to chronic neuroinflammation, synaptic loss, and dystrophic neurites. All studies were conducted following the NIH, American Physiological Society, and UCI Animal Care Guidelines. Mice were housed in cages containing cotton nestlet material with 4 or 5 animals per cage to minimize stress associated with social isolations. Any necessary veterinary care was provided by vivarium technicians, who check on mice daily for any changes in the health or well-being of the mice. Consistent with the recommendation of the Panel of Euthanasia of the AVMA, mice with continued discomfort and distress will be euthanized by inhalant overdose of CO_2_, followed by secondary euthanasia via cervical dislocation. All animal experiments involving mice were approved by UC Irvine Institutional Animal Care and Use Committee and conducted in compliance with all relevant ethical regulations for animal testing and research. All experiments involving mice also comply with the Animal Research: Reporting of *in vivo* Experiments (ARRIVE) guidelines.

### Viral infection

All infections were performed on mice under deep anesthesia through intraperitoneal (i.p.) injections of a mixture containing 85 mg/kg ketamine and 10 mg/kg xylazine. Subsequently, mice were infected intracranially (i.c.) with 500 plaque-forming units (PFU) of JHMV in 30µL of sterile Hanks balanced sterile solution (HBSS); control mice receive i.c. injections only containing HBSS. Infected and control mice were weighed and monitored daily using a well-accepted scoring criteria to screen clinical disease symptoms such as significant weight loss, inactivity, lack of grooming, and other indicators of distress or discomfort^26,27^.

### RNA Extraction

Brains were isolated at defined times 12 days post-infection (p.i.) was also homogenized with RNA extraction, cDNA synthesis, and qPCR for comparison of JHMV Membrane (M) mRNA levels performed as previously described^27^. Briefly, mouse brain tissue was added to TRIzol and homogenized using the Bead Ruptor 12 (Omni International) and 1.4mm ceramic beads (Omni International, 19-627). RNA was extracted via RNAeasy Minikit (Qiagen, 74106) using the “Purification of Total RNA, Including Small RNAs, from Animal Tissues” protocol from the manufacturer and Buffer RW1 to substitute Buffer RWT.

### cDNA synthesis and qPCR

cDNA was made from extracted RNA using previously described methods^28^. Briefly, cDNA was synthesized using the “First Strand cDNA Synthesis” protocol by New England Biolabs, using AMV Reverse Transcriptase (New England Biolabs, M0277L), Random Hexamers (Invitrogen, N8080127), RNAse Inhibitor (New England Biolabs M0314L) and AMV Buffer (New England Biolabs B0277A). Primer sequences used were GAPDH (forward: AACTTTGGCATTGTGGAAGG; reverse: GGATGCAGGGATGATGTTCT), JHMV Matrix glycoprotein (forward: TCAACCCCGAAACAAACAACC; reverse: GGCTGTTAGTGTATGG TAATCCTCA). qPCR was performed using the Bio-Rad iQ5 and iTaq TM Universal SYBR© Green Supermix (Bio-Rad, Hercules, CA). Reactions were performed using 10µL and the machine was set to run using the following parameters: 1 cycle (95°C for 3 minutes) followed by 40 cycles (95°C for 10 seconds, then 55°C for 30 seconds). Ct values for each sample were normalized to an internal control (GAPDH), yielding dCt values where lower dCt values indicated higher mRNA levels present while higher dCt values represented lower mRNA levels as more cycles of amplification was required to detect a signal across threshold.

### Cell Isolation and Flow Cytometry

Flow cytometry was performed to identify inflammatory cells entering the CNS using established protocols^27,29^. In brief, single cell suspensions were generated from tissue samples by grinding with frosted microscope slides. Immune cells were enriched via a 2-step Percoll cushion (90% and 63%), and cells were collected at the interface of the two Percoll layers. Before staining with fluorescent antibodies, isolated cells were incubated with anti-CD16/32 Fc block (BD Biosciences, San Jose, CA) at a 1:100 dilution. Immuno-phenotyping of cells was performed using commercially available antibodies specific for the following cell surface markers: CD4 (Invitrogen, 11-0042-82), CD8a (Invitrogen, 17-0081-82), CD11b (Abcam, ab24874) and CD45 (BioLegend, 103114; 103130). Cells were simultaneously incubated with LIVE/DEAD Aqua Dead Cell Stain (Invitrogen, L34966). The following flow cytometric gating strategies were employed for inflammatory cells isolated from the CNS: macrophages/myeloid cells (CD45 hi CD11b^+^) and microglia (CD45 lo CD11b^+^); FITC-conjugated rat anti-mouse CD4 and a PE-conjugated tetramer specific for the CD4 immunodominant epitope present within the JHMV matrix (M) glycoprotein, spanning amino acids 133-147 (M133-147 tetramer), to determine total and virus-specific CD4^+^ cells (37, 38); APC-conjugated rat anti-mouse CD8a and a PE-conjugated tetramer specific for the CD8 immunodominant epitope present in the spike (S) glycoprotein, spanning amino acids 510-518 (S510-518), to identify total and virus-specific CD8^+^ cells (37, 38). Data were collected using a Novocyte flow cytometer and analyzed with FlowJo software (Tree Star Inc.).

### Histopathology

6-month and 10-month-old 5xFAD hemizygous and wildtype non-transgenic mice were euthanized by inhalant overdose of Isoflurane at 7, 12, or 14-days post-infection before opening the thoracic cavity as a secondary method of euthanasia. Before transcardial perfusion with 1X phosphate buffered saline (PBS), blood plasma was collected from mice via cardiac puncture. In all experiments, brains were dissected, and the hemispheres were divided along the midline. One hemisphere of each brain was fixed in 4% paraformaldehyde (PFA) in PBS for 24 hours at 4°C for immunohistochemical and spatial single-cell transcriptomics. The other hemisphere was either fresh-frozen in dry ice for sample preparation to perform biochemical and protein analysis or placed in an enzyme digestion mix for cell isolation and staining to perform Flow cytometry. Additionally, spinal columns were isolated and stored in 4% PFA for 24-36 hours prior to analyzing spinal cord histopathology using Luxol Fast Blue staining protocols.

### Immunofluorescence Staining

After 24 hours, PFA-fixed brain hemispheres were transferred into 30% sucrose in 1X PBS for cryoprotection stored in 4°C for 48 - X hours before they were embedded in OCT and stored in −80°C. Sagittal brain sections were cut at 30µm using a cryostat (Thermo 95 664 OEC70 Micron HM525) and collected while free-floating in an anti-freeze solution of 30% ethylene glycol and 30% glycerol in 1X PBS kept in −20°C. Selected brain sections were chosen for immunohistological analyses between 1.10mm and 1.95 mm lateral to Bregma). One representative brain section from each mouse within the identical experimental cohort (i.e., same age, sex, genotype, and experimental condition) were subjected to simultaneous staining in the same container as described^24,30,31^. Briefly, the selected free-floating brain sections underwent several washes, at room temperature unless otherwise stated as follows: three 1X PBS washes for 5 minutes. For Amylo-Glo staining, free-floating brain sections were washed in 70% ethanol for 5 minutes followed by rinsing with deionized water for 2 minutes before immersed in Amylo-Glo RTD Amyloid Plaque Staining Reagent (1:100 dilution in 0.9% saline solution; TR-200-AG; Biosensis, Thebarton, South Australia) for 10 minutes per manufacturer’s instructions. Post-incubation, the free-floating sections were rinsed with 0.9% saline solution for 5 minutes before a brief wash in deionized water for 15 seconds. After Amylo-Glo staining, the brain sections continued onto standardized indirect immunohistochemical procedures as previously described^24,30,31^. Briefly, sections were immersed in blocking serum solution (5% normal goat serum with 0.2% Triton-X in 1X PBS) for 1 hour before an overnight incubation at 4°C in primary antibodies diluted in blocking serum solution. A complete list of antibodies with their respective dilutions is provided in Table 1. The following day, the brain sections were incubated with secondary antibodies for 1 hour in the dark after three washes of 1X PBS. Before mounting the brain sections onto microscope slides for imaging, the stained brain sections were once more rinsed in three washes of 1X PBS.

### Confocal Microscopy and Imaris Quantitative Analysis

Brain sections were imaged with a Zeiss Axio Scan Z1 Slidescanner using a 10X 0.45 NA Plan-Apo objective. High-resolution confocal fluorescence images were also obtained using a 20X 0.75 NA objective on a Leica TVS SPE-II confocal microscope. Two brain regions per mouse were selected for imaging: subiculum (SUB) and somatosensory cortex (SS CTX), unless otherwise stated and one field of view (FOV) per brain region was obtained for image analysis using Bitplane Imaris Software for quantification. Cell counting and volumetric measurements were acquired using the Spots and Surfaces modules through batch analysis of each imaged brain region.

### Protein Extraction, Biochemical Analysis via Meso Scale Discovery (MSD) Assays

Sample preparation and quantification of Aβ followed established protocols^30,31^. Hippocampal and cortical regions of each mouse brain hemisphere were micro-dissected and flash-frozen. Samples were pulverized using a Bessman Tissue Pulverizer. For Aβ biochemical analysis, pulverized hippocampal tissue was homogenized in 150µL of Tissue Protein Extraction Reagent (TPER; Life Technologies, Grand Island, NY). Cortical tissue was homogenized in 1000µL/150mg of TPER. The formulation of TPER contains 25mM bicine and 150 mM sodium chloride (pH 7.6) to effectively solubilize proteins within brain tissue post-homogenization. Protease (Roche) and phosphatase inhibitors (Sigma-Aldrich) were added to the homogenized samples and centrifuged at 100,000 g for 1 hour at 4°C to generate TPER-soluble fractions. To further create formic acid fractions, pellets from TPER-soluble fractions were homogenized in 70% formic acid: either 75µL or half of the added TPER volume for hippocampus or cortical tissue, respectively. Samples were once again centrifuged at 100,000 x *g* for 1 hour at 4°C. Proteins in this insoluble fraction were normalized to the respective brain region weight, while protein in the TPER-soluble fractions was normalized to the protein concentration determined via Bradford Protein Assay. Formic acid neutralization buffer (1 M TRIS base, 0.5 M Na_2_HPO_4_, 10% NaN_3_) was used to adjust pH before running ELISA assays. Quantitative biochemical analysis of human Aβ soluble and insoluble fraction levels were obtained using the V-PLEX Aβ Peptide Panel 1 (6E10) (K15200G-1; Meso Scale Discovery, Rockville, MD). Quantitative biochemical analysis of neurofilament-light chain (NfL) in plasma samples was performed using the R-Plex Human Neurofilament L Assay (K1517XR-2; Mesoscale Discovery).

### Luxol Fast Blue (LFB) Staining

Preparation for spinal cord histology was performed using previously described protocols^26,28^. In brief, PFA-fixed spinal columns were dissected to carefully extract the spinal cord from thoracic vertebrae 6-10 and cryoprotected in 30% sucrose for three days at 4°C. Spinal cords were then cut in 1mm transverse blocks and subsequently embedded in optimum cutting temperature (OCT) compound (VWR, Radnor, PA, USA) and frozen at −80°C. Spinal cord tissue was then coronally cryosectioned with a thickness of 8 micrometers (µm) and mounted on slides for luxol fast blue (LFB) staining using established protocols. Sections were stained with hematoxylin and eosin (H&E) in combination with LFB. Between 5–8 sections per mouse spinal cord were imaged using brightfield microscopy for analysis of demyelination. Areas of total white matter and demyelinated white matter were obtained using FIJI ImageJ Software, while demyelination was scored as a percentage of total white matter from the analyzed spinal cord sections as previously described.

### Bulk RNA Sequencing

RNA was extracted from 6-month-old 5xFAD mouse brains infected with 500 PFU JHMV at 7- and 14-days p.i. as described above. Library preparation, RNA sequencing, and read mapping analysis were performed by Novogene Co. Gene expression values were normalized into Log_2_ FPKM (fragments per kilobase of transcript per million mapped reads). Heatmaps were created using Morpheus (Morpheus, https://software.broadinstitute.org/morpheus). Volcano plots were created using custom code in R. Differentially expressed genes (DEGs) were filtered as significant with the magnitude of Log_2_ (Fold change) greater than 0.5 (|Log_2_FC| > 0.05) and a false discovery rate (FDR) < 0.05. For each experimentally relevant comparison, gene ontology (GO) term enrichment analysis was performed on significant DEGs using enrichR (https://amp.pharm.mssm.edu/Enrichr/).

### Spatial Transcriptomic Tissue Preparation

PFA-fixed brain hemispheres were embedded in OCT and stored frozen at −80°C. 24 hours prior to processing tissue for CosMx Spatial Molecular Imaging (SMI), sagittal brain sections were cut at 10µm and mounted on VWR Superfrost Plus microscope slides (Avantor, 48311-703). In each slide, six sagittal sections equally representing each experimental group were mounted and allowed to dry at room temperature for 15 minutes to promote tissue adherence and stored overnight at −80°C with desiccant. The following day, the brain tissue slides were processed (a total of 12 brain sections) in accordance with the Bruker Nanostring CosMx specifications. Unless otherwise noted, the following steps were performed at room temperature under a fume hood in a cleaned, RNAse free area. Slides were dried in an oven for 60 minutes at 60°C before briefly incubated in pre-cooled 10% neutral-buffered formalin for 15 minutes at 4°C and rinsed in three 1X PBS washes afterwards. The slides were placed back into the oven at 60°C for 45-60-minutes to further improve tissue adherence. After three washes in 1X PBS for 5 minutes, the slides were incubated in 4% sodium dodecyl sulfate (SDS) for 2 minutes, then washed in three washes of 1X PBS before dehydrating the tissue in subsequent 50%, 70% and 100% ethanol washes for 5 minutes each. After allowing the slides to air dry flat for 10-30 minutes, the slides were placed in a pre-heated container of 1X CosMx Target Retrieval Solution (Nanostring) at 100°C for 5 minutes, maintained by a steamer set to 100°C. After the antigen retrieval step, the slides were cooled in DEPC-treated water (ThermoFisher Scientific, AM9922) for 15 seconds and 100% ethanol for three minutes before 30 minutes of air drying. To permeabilize tissue, the sections were incubated in a digestion buffer (3 µg/mL Proteinase K in 1X PBS; Nanostring) for 30 minutes before a rinse in NBF Stop Buffer (0.1M Tris-Glycine Buffer, Cat# 15740) for 5 minutes, then rinsed in three 1X PBS washes for five minutes each. Meanwhile, a fiducial solution (Nanostring) was prepared to a 0.0015% dilution in 2X SSC-T and applied on tissue for 5 minutes. From this point, the slides were shielded from light. After two 1X PBS washes, slides were post-fixed in 10% NBF for five minutes and washed in two NBF Stop Buffer washes for five minutes each. Afterwards, the slides were left to incubate in an NHS-acetate (100mM; Cat#26777) solution for 15 minutes before being rinsed in 2X saline-sodium citrate (SSC, Cat#AM9763) for 5 minutes. Finally, the tissue was left to incubate with probe mix containing segmentation markers for ribosomal RNA (rRNA), CosMx Mouse Neuroscience Panel, 1000-plex, RNA with a custom RNA Add-On set, and RNAse inhibitor in Buffer R (Nanostring). The slides were placed inside a staining tray and left inside oven set to 37°C for hybridization over 16-18 hours overnight at which point, the hybridization probe mix was removed from the slides, and these were placed in two pre-heated stringent washes (50% deionized formamide, CAT #AM9342) in 2X SSC at 37°C for 25 minutes each. After two washes in 2X SSC for two minutes, the tissue was incubated in DAPI nuclear stain for 15 minutes and then washed in 1X PBS for 5 minutes. Meanwhile, a segmentation marker mix for GFAP and histones was prepared and left to incubate on the tissue for 60 minutes. After slides were rinsed in three 1X PBS washes for 5 minutes, the tissue was once again incubated in NHS-Acetate before finally being rinsed in two 2X SSC washes for 5 minutes. Before loading into the CosMx Spatial Molecular Imaging instrument, adhesive flow cells (Nanostring) were placed in each slide to create a fluidic chamber for imaging according to manufacturer’s instructions. Once loaded and processed in the machine according to the manufacturer’s instructions for a 2-slide run, approximately 850 fields of view (FOVs, about 75 FOVs per brain section) was chosen to capture the following brain regions: cortical regions, corpus callosum, hippocampus, upper thalamus, and upper caudate striatal regions for each of the 12 brain sections. Imaging continued for approximately 6 days and the raw data was collected onto the Nanostring AtoMx online platform.

### Spatial Transcriptomic Analysis

For spatial transcriptomic analysis, raw datasets were exported from Nanostring AtoMx as a Seurat object and processed with R 4.3.1 software as previously described^31,32^. Briefly, principal component analysis (PCA) and uniform manifold approximation and projection (UMAP) analysis was performed to reduce dataset dimensionality and visualize cell clustering at 1.0 resolution to yield 42 clusters. Clusters were manually annotated based on top transcript expression markers of known marker genes and its location in XY space. Furthermore, cell count and proportion plots were generated by plotting the number of cells in each cell type and scaling values relative to 1) normalized percentages per experimental group by obtaining the ratio of cell counts in each cell type-group pair by the total number of cells and 2) dividing by the sum of the proportions across the cell type to account for differences in sample sizes. MAST was used on scaled expression data to enable differential gene expression analysis per cell type between group comparisons to compute the average difference as defined by the difference in log-scaled average expression across the compared groups for each cell type. DEG scores were calculated by take the sum of the absolute log_2_ fold change values of all transcripts with statistically significant (p_adj_ < 0.05) differential gene expression patterns between two group comparisons for each cluster. To generate data visualization, we utilized ggplot2 3.4.4174.

### Statistics

Sample sizes were calculated using power analyses based on pilot data (G*power parameters: power 0.95, Type I Error Rate 5%, Effect Size: 1.56) and on historical experience (i.e., mice lost due to acute infection or aging), while being powered to detect relevant sex differences. Animal groups are randomized at weaning, and age-matched control cohorts will be generated alongside their respective experimental groups to maximize rigor and reproducibility. Data was analyzed using two-tailed unpaired t-tests for comparison between two groups or ANOVA (one- or two-way with mixed effects) for comparisons of up to four groups with appropriate post-hoc tests (Holm-Šidák’s multiple comparisons) when necessary. For non-parametric data, Kruskal-Wallis tests were used to compare group differences followed by Dunn’s multiple comparisons test. Statistical significance is defined as p<0.05 and statistical trends at p<0.1. Read mapping analysis and statistics for bulk RNA sequencing data were performed by Novogene Co. Statistics for transcriptomic data (CosMx Spatial Molecular Imager) will be calculated using standard R packages, with a False Discovery Rate (FDR) <0.05.

## Results

### 5xFAD mice exhibit similar weight loss, mild clinical disease, and control of viral replication within the CNS following intracranial JHMV infection

To determine whether familial AD mutations and its ongoing neuroinflammation impacted morbidity following JHMV infection of the CNS, we utilized 10-month-old 5xFAD and wildtype (WT) C57BL/6 mice to measure viral replication and onset of neurological disease during acute JHMV-mediated encephalitis at 12 days post-infection (p.i) **(Figure 1A)**. By 12 days p.i., both JHMV-infected WT mice and JHMV-infected 5xFAD mice had significant reductions in body weight compared to uninfected controls (**Figures 1B&C**, p<0.0001 and p<0.01, respectively). When comparing the infected groups together, JHMV-infected wildtype mice had greater body weight loss compared to JHMV-infected 5xFAD mice (p<0.05). When considering clinical disease severity assessed by hindlimb paralysis, both JHMV-infected WT and JHMV-infected 5xFAD mice displayed similar progression of motor impairment and clinical disease by 12 days p.i. **(Figure 1D)**. By 12 days p.i., both JHMV-infected WT and JHMV-infected 5xFAD mice had significant progressive motor impairments compared to their uninfected controls (**Figure 1E**, p<0.0001 and p<0.05, respectively). Using qPCR to quantify levels of viral RNA in infected brains by 12 days p.i., no significant differences were observed between JHMV-infected WT and JHMV-infected 5xFAD, suggesting similar control over viral replication at 12 days p.i. **(Figure 1F)**. This is corroborated in younger 6-month-old WT and 5xFAD mice, which also displayed similar levels of viral RNA via qPCR by 10-14 days p.i. **(Supplementary Figure 1A)**.

**FIGURE 1.**
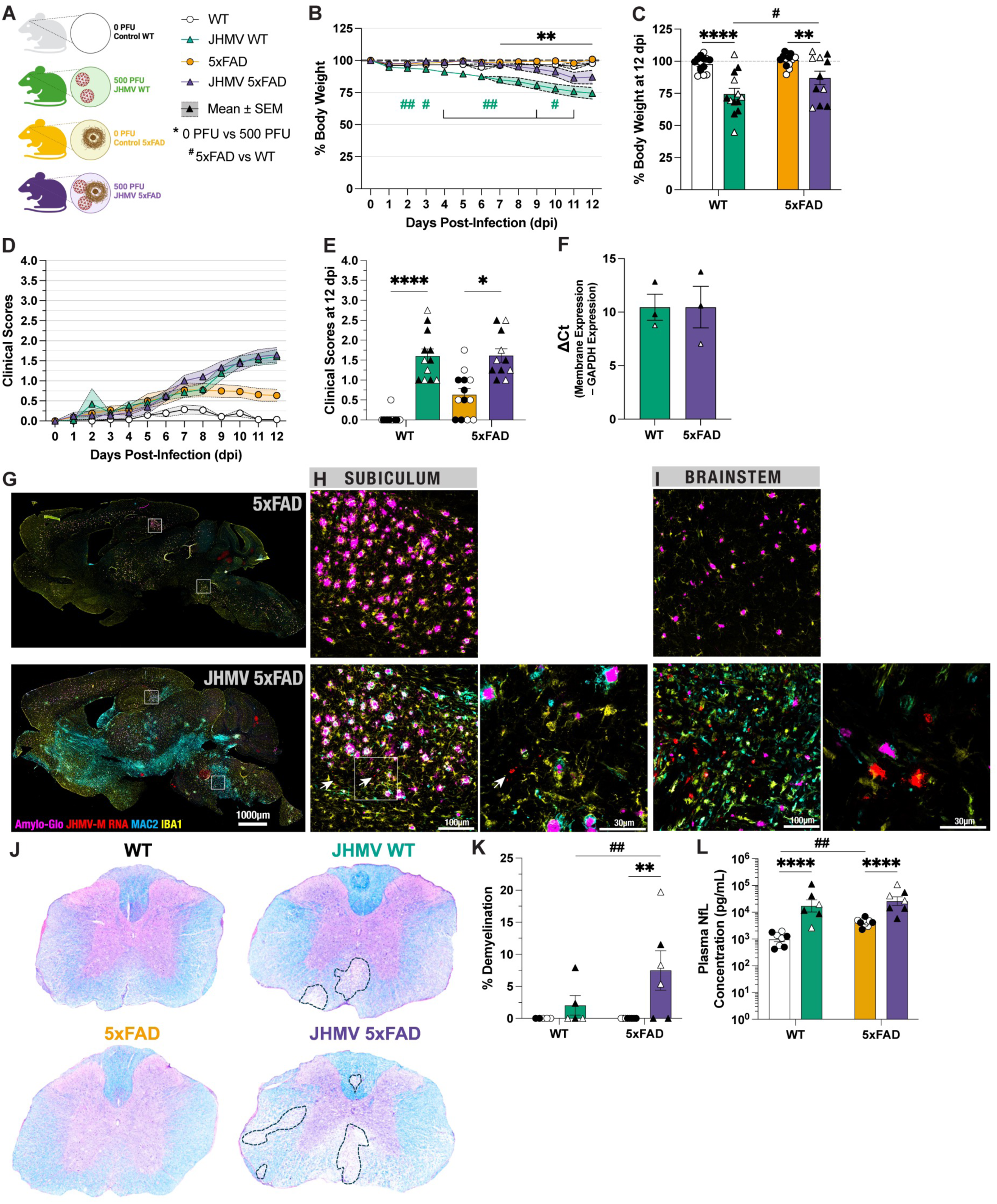
JHMV infection leads to reduced body weights, increased clinical disease severity, and demyelination in spinal cords despite similar levels of viral control in infected brains at 12 dpi. **A)** Schematic of each experimental group. **B)** Line chart depicting changes in initial body weights across each day post-infection for uninfected WT (white), JHMV-infected WT (green), uninfected 5xFAD controls (yellow), and JHMV-infected 5xAFD mice (purple). **C)** At 12 dpi, percent changes in body weight were compared across each experimental group. **D)** Line chart depicting the progression of clinical disease severity and hindlimb paralysis across each day post-infection. **E)** At 12 dpi, clinical disease scores were compared across each experimental group. **F)** qPCR with extracted RNA from brain homogenates demonstrate similar transcript levels of viral membrane protein RNA between JHMV-infected WT and 5xFAD brains at 12 dpi. **G)** Representative whole-brain scanned images of sagittal brain sections from uninfected and JHMV-infected 5xFAD brains co-stained with Amylo-Glo, MAC2, and IBA1 in addition to *in situ hybridization* targeting JHMV RNA. Representative 20X and 63X confocal images of subiculum **(H)** and brainstem regions **(I)** in brain sections of uninfected and JHMV-infected 5xFAD mice. Fluorescent signals of viral RNA are indicated with white arrows. **J)** Representative images of H&E/Luxol Fast Blue-stained spinal cord sections across each experimental group. Dashed lines demarcate areas of demyelination. **K)** The percentage of demyelination was compared across each experimental group. **L)** Plasma concentration of NfL protein in blood plasma was compared across each experimental group. n=7-10 per group. Data is presented as mean ± SEM. Unpaired t-tests was used when appropriate and two-way ANOVA followed-by Holm-Sidak post-hoc tests were performed to examine biological relevant interactions. Clinical disease scores were compared using Kruskal-Wallis tests followed by Dunn’s multiple comparisons test. *p<0.05, **p<0.01, ***p<0.001, ****p<0.0001. Males are represented with closed symbols and females are represented with open symbols.

### JHMV viral RNA are present across similar brain regions in the CNS of infected WT and 5xFAD mice

Despite effective viral clearance, sterile immunity is not achieved and viral antigen and RNA will continue to persist within white matter of the CNS. Using antibodies against JHMV nucleocapsid protein, histological staining revealed presence of viral antigen within white matter regions (e.g., brainstem and peduncles) in addition to the subiculum hippocampal region within JHMV-infected 5xFAD brains at 12 days p.i. (Supp. Figure Figure 1B-C). Notably, viral antigen is not observed in somatosensory cortical regions. In addition to viral antigen, in situ hybridization (ISH) via RNAscope also reveals the widespread active replication of viral RNA throughout the wildtype brain at 7 days p.i. within specific regions where viral antigen is present, specifically the subiculum and brainstem (Supp. Figure Figure 1D-F). This is also corroborated in JHMV-infected 5xFAD brains, which reveal a lingering presence of JHMV RNA in the subiculum region, but more so in the brainstem (**Figure 1G-I**). Despite similar viral control efficacy, JHMV-infected 5xFAD mice exhibited greater immune-mediated demyelination in Luxol-Fast Blue (LFB)-stained spinal cord sections at 12 days p.i. compared to JHMV-infected WT mice (**Figure 1J-K**, p<0.01). To assess levels of neuronal damage resulting from JHMV infection, we measured concentration of plasma neurofilament light chain (NfL). In line with previous studies, we found significant increases in plasma NfL protein concentrations in 5xFAD mice compared to WT mice **(Figure 1L)**. Additionally, we found that JHMV infection further increased plasma NfL concentration in JHMV-infected WT and 5xFAD compared to their respective, uninfected controls **(Figure 1L)**.

### Dense-core Aβ plaque burden is reduced in JHMV-infected 5xFAD mice in subiculum and somatosensory cortical regions

The significance of viral RNA present in the subiculum compared to the somatosensory cortex may suggest that areas of viral RNA persistence may influence Aβ plaque deposition in 5xFAD mice^24,33^. Staining for OC^+^ fibrillar Aβ surrounding the denser plaques revealed no changes in either 5xFAD subiculum or somatosensory cortex regions following JHMV infection (**Figures 2A, C-D**). Staining for Aβ_1-16_ residues with 6E10 demonstrated similar results, with only trending reductions in 6E10^+^ plaque volume within the subiculum following JHMV infection (**Figure 2B, E-F**). This is further corroborated by biochemical analysis demonstrating no differences in soluble and insoluble Aβ40 and Aβ42 protein concentration in cortex homogenates of JHMV-infected and control 5xFAD brains via MULTI-ARRAY assay (Mesoscale Discovery) (**Figure 2G-H**). However, Amylo-Glo staining for dense-core Aβ plaques revealed significant reductions in Aβ plaque volume in the subiculum (**Figure 3A,B,D-F**, p<0.01). However, within the somatosensory cortex, where viral RNA was not observed, JHMV infection significantly reduced the number of dense-core Aβ plaques with only a trending reduction in the average volume of Amylo-Glo^+^ dense-core plaques (**Figures 3A,C, G-I**, p<0.0001). These results indicate that JHMV infection and its resulting encephalitis may distinctly impact dense-core plaques as opposed to overall Aβ burden in the 5xFAD mouse brain.

**FIGURE 2.**
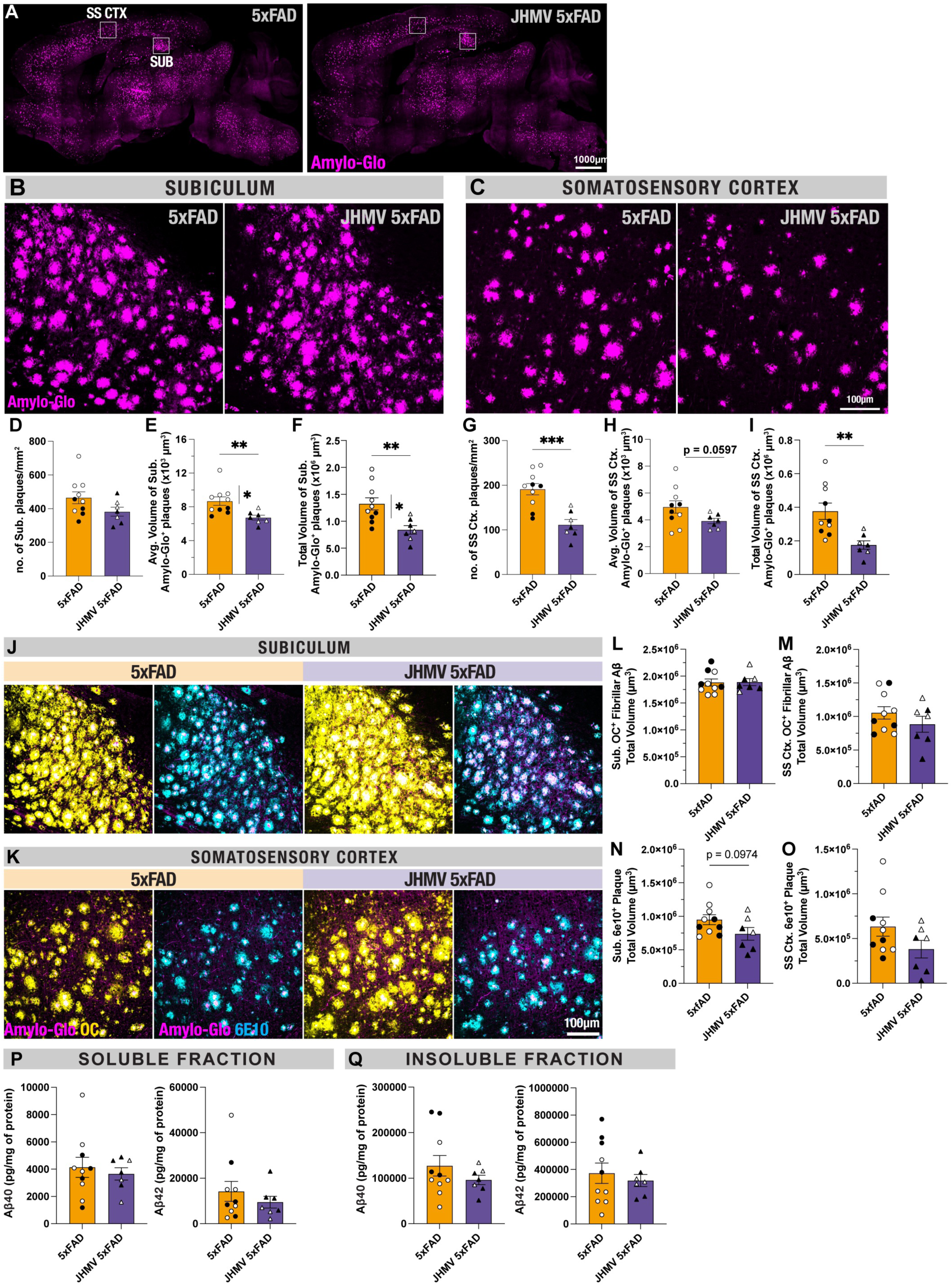
JHMV infection reduces dense-core amyloid plaque pathology, but not overall Aβ burden. Aβ plaque burden was assessed using Amylo-Glo staining for dense-core plaques in 30µm sagittal brain sections. **A)** Representative whole-brain images of 12dpi JHMV-infected and control 5xFAD mouse brains stained with Amylo-Glo. Representative 20X confocal images of subiculum **(B)** and somatosensory cortex **(C)** depicting Amylo-Glo^+^, dense-core Aβ plaques. Quantification of plaque densities **(D, G)**, individual plaque volumes **(E, H)**, and average plaque volumes **(F, I)** with the field-of-view (FOV) of JHMV-infected and control 5xFAD brain sections. Representative confocal images of brain sections costained with Amylo-Glo, conformation-specific OC fibrils, and Aβ_1-16_ residues in subiculum **(J)** and somatosensory cortex **(K)** of uninfected and JHMV-infected 5xFAD mice. Quantification of OC fibrillar Aβ volumes around dense-core plaques in subiculum **(L)** and somatosensory cortex **(M)**. Quantification of 6e10^+^ plaque volumes in subiculum **(N)** and somatosensory cortex **(O)**. Protein concentration of Aβ40 and Aβ42 within detergent-soluble fractions **(P)** or insoluble fractions **(Q)** of cortex homogenates. of n=7-10 per group. Data is presented as mean ± SEM. Unpaired t-tests was used to examine statistically significant differences between groups. *p<0.05, **p<0.01, ***p<0.001, ****p<0.0001. Males are represented with closed symbols and females are represented with open symbols

**FIGURE 3.**
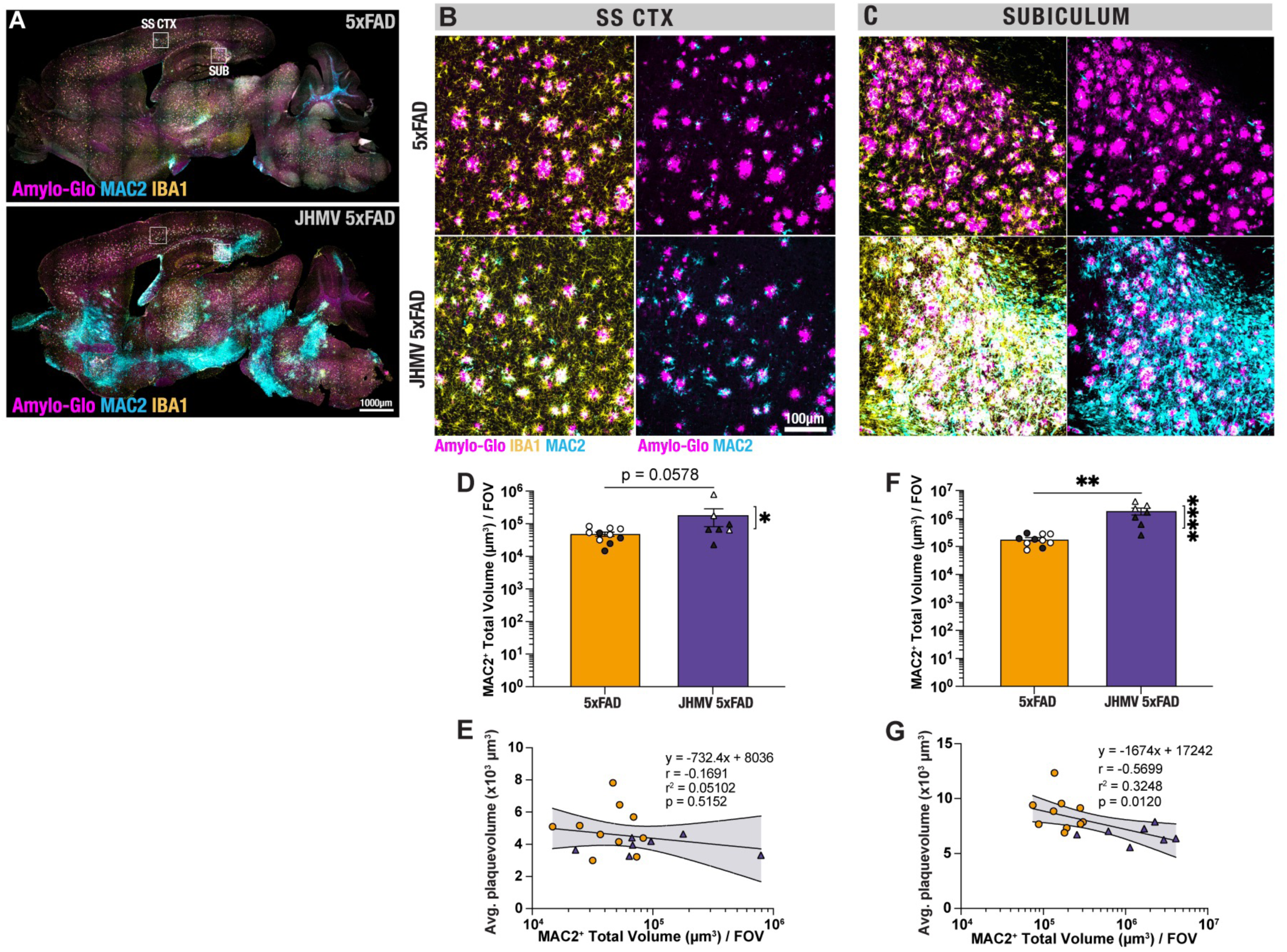
Regions of high infiltration of MAC2^+^ peripheral myeloid cells are associated with reduced Aβ plaque volumes in JHMV-infected 5xFAD mice. Infiltration of peripheral myeloid cells (i.e., bone-marrow-derived monocytes and monocyte-derived macrophages) was assessed using MAC2 antibody immunostaining of 30µm sagittal brain sections. **A)** Representative whole-brain images of 12dpi JHMV-infected and control 5xFAD mouse brains co-stained with Amylo-Glo, MAC2, and IBA1. Representative 20X confocal images of somatosensory cortex (**B)** and subiculum**(C)** depicting Amylo-Glo^+^ dense-core Aβ plaques surrounded by MAC2^+^ and IBA1^+^ macrophages. Quantification for total volume of MAC2^+^ cells in somatosensory cortex **(D)** or subiculum **(F)** of JHMV-infected and control 5xFAD brain sections. Scatterplot and linear regression depicting no relationship in somatosensory cortex **(E)**, yet a significant correlation between total volume of MAC2^+^ cells and average plaque volumes in subiculum **(G)** 4in 5xFAD mice. n=7-10 per group. Data is presented as mean ± SEM. Unpaired t-tests was used to examine statistically significant differences between groups. Spearman correlation and linear regression was performed to measure the relationship between MAC2^+^ cell volume and Aβ plaque volume. **p<0.01. Males are represented with closed symbols and females are represented with open symbols.

### MAC2^+^ macrophages infiltrate into the CNS following JHMV infection and strongly correlate with brain regions demonstrating reduced Aβ plaque burden

We have previously demonstrated that inflammatory *Lgals3*/MAC2-expressing monocyte/macrophages migrate to areas of viral replication in response to chemokine expression using a combination of single-cell RNA sequencing and histology^34^. Immunostaining *Lgals3/*MAC2 in JHMV-infected 5xFAD brains demonstrated high infiltration of peripheral monocyte/macrophages in the same brain regions where JHMV viral antigen is present, namely the subiculum, peduncles, and brainstem (**Figure 4A**). Representative confocal images of AD-relevant brain regions like the somatosensory cortex and subiculum also illustrated localized interactions between MAC2^+^ and Aβ plaques (**Figure 4B-C**). Moreover, quantifying the activation state MAC2^+^ cells via assessment of cell volume within AD-relevant regions revealed increased volumes of MAC2^+^ cells within the somatosensory cortex (p<0.0578) and a significant increase (p<0.01) in volume of these cells in the subiculum (**Figures 4D&F**). Moreover, there was a significant (p<0.05) correlation between enriched MAC2^+^ infiltration within the subiculum and associated with reduced Aβ plaque volumes (**Figure 4G**, r^2^=0.3248, p=0.0120). These findings suggest that inflammatory monocyte/macrophages that are recruited into the CNS in response to JHMV replication during acute disease contribute to reduced morphology of dense-core Aβ plaques.

**FIGURE 4.**
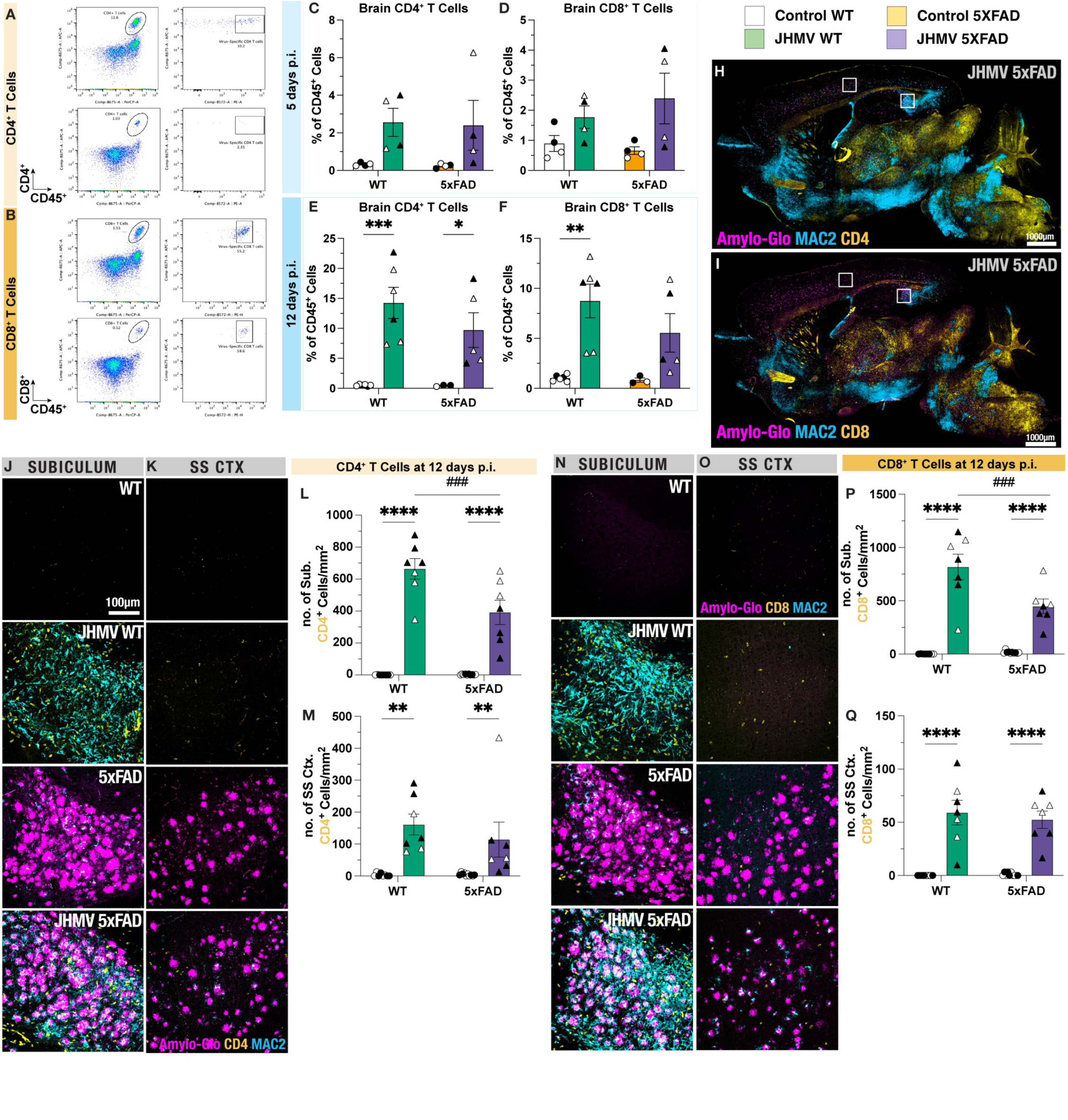
T cells infiltrate into the brains of both JHMV-infected WT and 5xFAD mice by 12 dpi into subiculum brain regions. Gating strategy to select CD45^+^ CD4^+^ T cells **(A)** and CD45^+^ CD8^+^ T cells **(B)** following cell isolation, staining, and flow cytometry. Flow cytometric analysis of CD4^+^ T cell frequency **(C)** and CD8^+^ T cell frequency **(D)** from brain isolates at 5 dpi. Flow cytometric analysis of CD4^+^ T cell frequency **(F)** and CD8^+^ T cell frequency **(F)** from brain isolates at 12 dpi. Infiltration of CD4^+^ and CD8^+^ T cells into brain regions were assessed with brain sections co-stained with Amylo-Glo, MAC2, and either CD4 **(H)** or CD8 **(I)** antibody. Representative 20X confocal images of subiculum **(J)** and somatosensory cortex **(K)** depicting the presence of CD4^+^ T cells within brain sections from each experimental group. Quantification of CD4^+^ densities per FOV in subiculum **(L)** and somatosensory cortex **(M)** and within each experimental group. Representative 20X confocal images of subiculum **(N)** and somatosensory cortex **(O)** depicting the presence of CD8^+^ T cells within brain sections from each experimental group. Quantification of CD8^+^ densities per FOV in subiculum **(P)** and somatosensory cortex **(Q)** within each experimental group. n=4-10 per group. Data is presented as mean ± SEM. Two-way ANOVA followed-by Holm-Sidak post-hoc tests was performed to examine biological relevant interactions. *p<0.05, **p<0.01, ***p<0.001, ****p<0.0001. Males are represented with closed symbols and females are represented with open symbols.

### CD4^+^ and CD8^+^ T cells infiltrate into the CNS following JHMV infection and tightly associate with MAC2^+^ interactions with Aβ plaques

To address the potential involvement of CNS inflammatory CD4^+^ and CD8^+^ T cells during acute JHMV infection in 5xFAD mice, flow cytometry was performed to quantify the frequency of CD45^+^ CD4^+^ and CD45^+^ CD8^+^ T cells at 5 and 12 days p.i. (**Figure 5A-B**). By 5 days p.i., there was moderate infiltration of both CD4^+^ and CD8^+^ T cells in both JHMV-infected WT and 5xFAD brains, although no significance was detected (**Figure 5C-D**). However, both JHMV-infected WT and 5xFAD brains exhibited significant levels of infiltrating CD4^+^ T cells at 12 days following JHMV infection compared to uninfected controls (**Figure 5E**, p<0.001 and p<0.05, respectively). Interestingly, JHMV-infected 5xFAD brains showed a trending increase in infiltrating CD8^+^ T cells despite JHMV-infected WT mice exhibiting a greater significant infiltration of CD8^+^ T cells compared to uninfected WT controls (**Figure 5F**, p<0.01).

**FIGURE 5.**
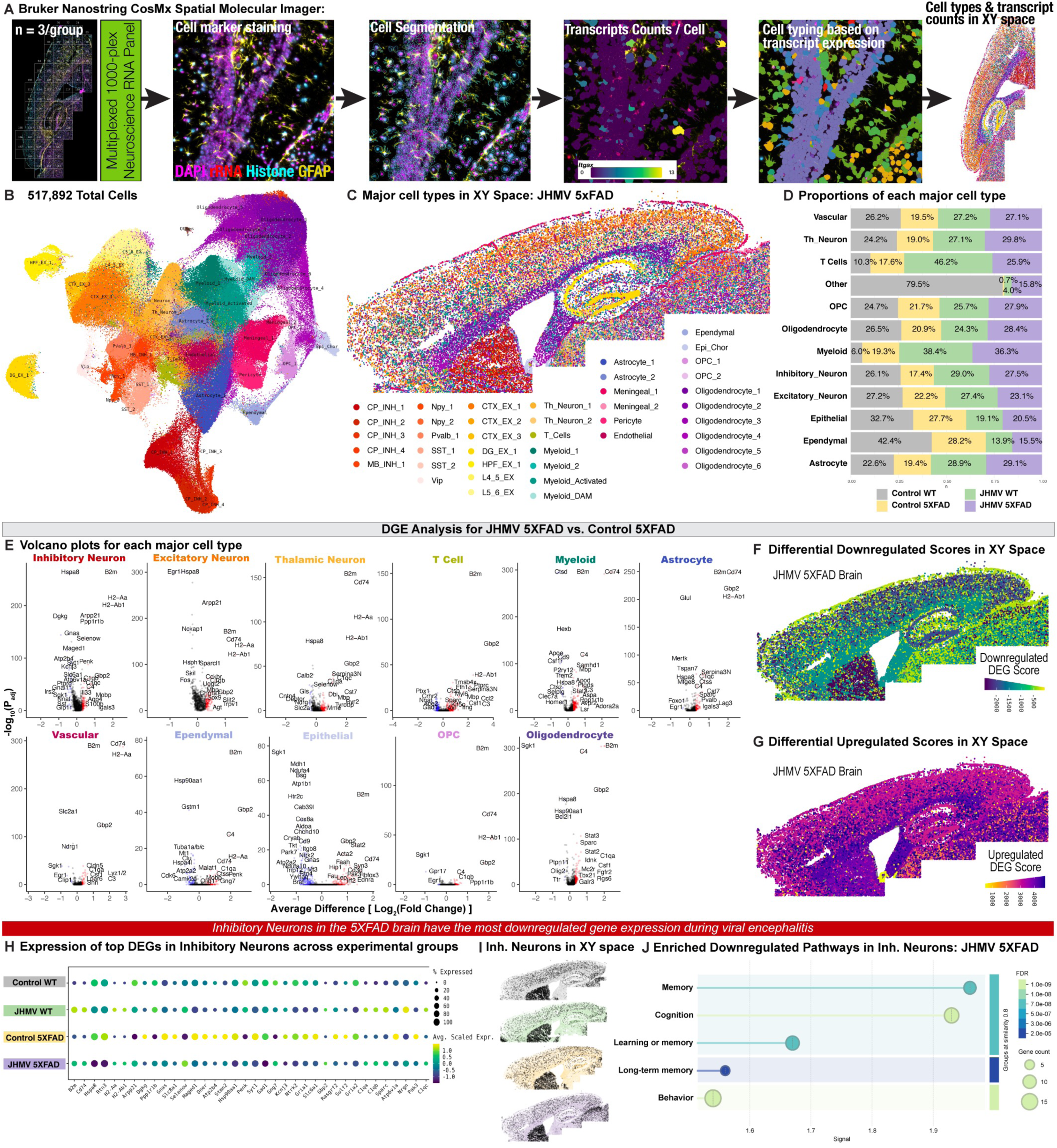
Spatial transcriptomic analysis of JHMV-infected and control WT and 5xFAD mice. **A)** Experimental workflow for targeted 1000-plex single cell spatial transcriptomic imaging of 12 sagittal mouse brain sections (n=3/group). Fields of view (FOVs) was selected in dentate gyrus to represent the cell marker staining, segmentation, transcript counts, and cell typing based on transcript expression. DAPI, rRNA, Histone, and GFAP were used as cell markers. **B)** Uniform Manifold Approximation and Projection (UMAP) of 517,892 cells across all brain samples were captured with a mean transcript count of 841 transcripts per cell. Unbiased cell clustering at 1.0 resolution identified 42 clusters, which were manually annotated with a combination of automated and manual approaches with reference to Allen Brain Atlas single-cell RNA-seq cell types, gene expression, and anatomic location in XY space. **C)** 42 annotated clusters plotted in XY space with a representative JHMV-infected 5xFAD brain. **D)** Proportions of each major CNS cell type grouped by experimental group. **E)** Volcano plots of DEGs within each major CNS cell type across JHMV-infected 5xFAD vs. control 5xFAD. **F)** Differential downregulation (DD) and **G)** differential upregulation (DU) scores for JHMV-infected 5xFAD vs. control 5xFAD in each cluster plotted in XY space using a representative JHMV-infected 5xFAD brain. **H)** Pseudo-bulk expression of top DEGs in inhibitory neurons between JHMV-infected 5xFAD vs. control 5xFAD, grouped by experimental group. **I)** Inhibitory neurons in XY space in representative brain sections from each experimental group. **J)** Pathways enriched via Gene Ontology (GO) of downregulated DEGs in JHMV-infected 5xFAD vs. control.

To visualize the infiltration of T cells into the brain parenchyma, we utilized fluorescent immunostaining of CD4^+^ and CD8^+^ T cells in tissue sections for confocal microscopy to also determine T cell populations in regions where reduced Aβ pathology was observed following viral encephalitis (**Figures 5H-I**). In parallel with the obtained flow cytometric data, JHMV-infection mediated the infiltration of effector T cells, resulting in significant increases in the density of CD4^+^ T cells in both subiculum and somatosensory cortex regions of infected WT and 5xFAD brains at 12 days p.i. (**Figure 5J-M**, p<0.0001 and p<0.01, respectively). Furthermore, immunostaining for CD8^+^ T cells also reveals increased infiltration of CD8^+^ T cells in subiculum and somatosensory regions within infected WT and 5xFAD brains (**Figure 5 N-Q**, p<0.0001). Interestingly, the infiltration of CD4^+^ and CD8^+^ T cells was significantly diminished in JHMV-infected 5xFAD mice compared to JHMV-infected WT mice (**Figure 5L and 5Q**, p<0.001).

### Bulk RNA sequencing reveals upregulation of genes associated to host immunity and viral-induced neuroinflammation in JHMV-infected mice

To investigate transcriptional changes within 5xFAD brains following viral encephalitis, we isolated RNA from frozen hemi-brain samples from each experimental group at days 7 and 14 p.i. and performed bulk sequencing analysis for dysregulated or differentially expressed genes (DEGs) for pathway analysis by gene ontology at each timepoint. Volcano plots were plotted to compare the average difference in gene expression (shown as Log_2_ fold change) in comparison with its respective control group. Genes were determined to be significantly dysregulated if the magnitude of average difference from baseline was greater than 0.5 with a false discovery rate/adjusted p-value less than 0.05 (i.e. |Log_2_FC| > 0.5, FDR > 0.05). Significant DEGs were then plotted on volcano plots and heatmaps to illustrate its level of differential expression. JHMV-induced encephalitis at both day 7 and 14 p.i. resulted in significantly down-regulated DEGs in JHMV-infected 5xFAD brains include *Sqle, Msmo1, Hmgcs1,* and *Gm9946.* Meanwhile, significantly DEGs in JHMV-infected 5xFAD brains such as *Lyz2, Cd68, Gpnmb, Ctss, Ctsb, Ctsd, Lgals3, Il7r, Ly6c2,* and *Cxcl9* were significantly up-regulated at day 7 p.i. **(Supplementary Figure 2A&J)**. Heatmap plots of the JHMV 5xFAD vs. Control 5xFAD comparison depict the magnitude of significant DEGs resulting from viral encephalitis **(Supplementary Figure 2B&K)**. Using Gene Ontology (GO), we conducted pathway analysis for down-regulated pathways enriched in DEGs impacted by viral encephalitis in the 5xFAD brain and observed enriched pathways associated with cholesterol and steroid biosynthesis, neuronal transmission, axonogenesis, and lipid and lipoprotein metabolism. As expected, viral encephalitis in 5xFAD brains also up-regulated pathways involved in immunity, such as the IL-2 signaling, cytokine signaling, T cell regulation of apoptosis, neutrophil activation, and lysosome functioning **(Supplementary Figure 2C&L)**. By day 14 p.i., several of these DEGs and enriched pathways impacted by viral encephalitis continue to be up-regulated, in addition to *Spp1, Ccl5,* and *B2m* **(Supplementary Figure 2J-L)**.

To determine whether these changes were specifically associated with JHMV-induced encephalitis, we have also performed bulk sequencing and DEG analysis on hemibrains from non-transgenic wildtype mice infected with JHMV at 7 and 14 days p.i. **(Supplementary Figures 2D-F & 2M-O)**. Indeed, many significant DEGs and enriched pathways were also shared in JHMV-infected wildtype mouse brains at 7 **(Supplementary Figures 2D-F)** and 14 days p.i. **(Supplementary Figures 3M-O)**. Comparing gene expression differences between JHMV 5xFAD vs. JHMV wildtype mouse brains minimal differences in genes associated with immunity at day 7, yet several significant genes appeared up-regulated in JHMV-infected 5xFAD brains such as *Plin4, Hif3a, Chil3, Cdc25c, Fam107a, Gm29650,* and *Kdm5d*. Gene ontology analysis of significant DEGs in JHMV 5xFAD vs. JHMV wildtype brains reveal enriched down-regulated pathways involved in BDNF signaling, while enriched up-regulated pathways include hemostasis, platelet activation, IL-6 regulation, and blood-brain-barrier transport **(Supplementary Figures 2G-I & 2P-R)**.

### Utilizing spatial transcriptomics to explore the impact of acute viral encephalitis on CNS cell types

While these bulk sequencing methods have identified key biological pathways impacted by viral encephalitis in the 5xFAD brain, we lack the spatial and single-cell resolution to determine which key cell types within the CNS are mediating changes associated with Aβ pathology. To determine the impact of acute viral encephalitis following JHMV infection on cells within the mouse brain we investigated transcriptional changes to myeloid cells and other CNS cell types using spatial transcriptomic imaging. This approach facilitates gene expression analysis with single-cell resolution of CNS cell types, previously limited by technique-induced expression changes associated with single-cell and nucleus RNA-seq^31^. Furthermore, this technique retains gene expression of cells in distinct spatial regions of the brain, enabling *in situ* quantification and profiling of cell populations within a brain slice. We performed this experiment utilizing a high-plex *in situ* analysis platform (Bruker Nanostring CosMx Spatial Molecular Imager) with a 950-plex RNA mouse neuroscience panel and a 50-plex Custom Add-On RNA panel (Bruker Nanostring)^31^. Three sagittal hemibrain slices per group were processed capturing the cortex and hippocampus regions, resulting in a dataset comprising of 517,892 cells from 12 brain sections. The CosMx SMI platform performed cell segmentation using DAPI, histone, ribosomal RNA, and GFAP staining **(Figure 5A, Supplementary Figure 3A)** and transcript counts per cell for each of the 1000 gene targets per cell were calculated with an average of 841 transcripts per cell for 517,892 cells **(Figure 5B, Supplementary Figures 3B&C)**. Cell clustering was obtained using a community detection approach on a k-nearest neighbor graph, followed by dimensionality reduction via Uniform Manifold Approximation and Projection (UMAP). Clusters were manually annotated based on top gene expression, expression of canonical CNS cell markers, and its spatial XY location in the brain, identifying 42 annotated clusters comprised of 11 clusters of inhibitory neurons, 7 clusters of excitatory neurons, 2 clusters of thalamic neurons, 2 clusters of astrocytes, 4 clusters of endothelial and vascular-related cell types, 4 clusters of myeloid cells, 2 clusters of oligodendrocyte precursor cells (OPCs), 6 clusters of oligodendrocytes, 1 cluster of T cells, 1 cluster of ependymal cells, 1 cluster of epithelial cells, and 1 cluster designated “other” **(Figure 5B&C, Supplementary Figures 3D&E)**. Each of the 42 clusters were condensed into major clusters comprised of similar cell types (e.g., Astrocyte_1 and Astrocyte_2 were combined into a broader “Astrocyte” cluster) **(Supplementary Figure 3F)**. The proportions of each major cell type cluster were then plotted per experimental group to illustrate any changes associated with JHMV-induced encephalitis, Aβ pathology, or both **(Figure 5D, Supplementary Figure 3F)**. Since cell clustering relies purely on gene expression levels and not spatial location, we verified the correct assignment of cell populations in neuroanatomical locations by assessing cell type proportions across each experimental group and by plotting cells in XY space **(Supplementary Figures 4A&B)**. For each cluster, we also determined cell counts across each cluster of cell types within the CNS **(Supplementary Figures 4C&D)**. Clusters with fewer than 500 cells (i.e., “Other”) were removed from downstream analysis. In concordance with histology and flow cytometry, we observed increased proportion of cells in myeloid and T cell clusters following JHMV encephalitis **(Supplementary Figures 4C&D)**.

To validate the impact of acute viral encephalitis on the transcriptional state of various cell types within the brain, we next conducted differential gene expression (DGE) analysis on JHMV-infected WT and control, uninfected WT. Volcano plots depict several changes in gene expression within each CNS subtype and major cell type in the brains of these two groups **(Supplementary Figure 5 & 6A)**. To visualize the spatial distribution of all cell types with changes in gene expression, we generated a DEG score representing the magnitude of fold change between each comparison of two experimental groups. This was calculated using the sum of products between two variables: the average difference in gene expression between two groups and the negative log 10 p-value for each gene^31^. The calculated value provides each cluster with a value plotted in XY space, depicting the DEG score of all differentially down-regulated (DD score) or up-regulated genes (DU score) **(Supplementary Figures 6B-C)**. Notably, the myeloid cell cluster demonstrated the greatest DD score within JHMV-infected WT brains, indicating the greatest down-regulation of genes in this cell population. Visualizing the location of myeloid cells in XY space on a JHMV-infected WT brain demonstrates that several myeloid cells exhibiting the most down-regulation are found throughout cortical regions and the subiculum of the hippocampus, as well as the fornix **(Supplementary Figure 6B)**. Pathway analysis of down-regulated DEGs within myeloid cells illustrate impacts on neuron projection morphogenesis, behavior, regulation of tissue and bone remodeling, and modulation of synaptic transmission **(Supplementary Figure 6D)**. Furthermore, myeloid cell populations significantly upregulate expression of several DEGs involved in gliogenesis, lymphocyte immunity, immune effector processes, and antigen-presentation consistent with previous single-cell sequencing analyses on JHMV infection of the CNS **(Supplementary Figure 6E)**.

In addition to myeloid cells also exhibiting high DU scores in JHMV WT brains, astrocytes and oligodendrocytes demonstrate the greatest magnitude of up-regulated genes during acute viral encephalitis **(Supplementary Figure 6A)**. Indeed, visualizing DU scores of each CNS cell in XY space illustrates the highest scores along white matter areas such as the corpus callosum and fornix, but also distributed throughout the cortex and hippocampal formation **(Supplementary Figure 6C)**. Pathway analysis using gene ontology (GO) of significantly up-regulated genes within astrocytes and oligodendrocytes reveal biological processes associated with immune responses, immune effector processes, gliogenesis, and synaptic pruning during acute viral encephalitis **(Supplementary Figures 6F&G)**.

### Spatial transcriptomic imaging reveals a shift in disease-associated myeloid cells surrounding Aβ plaque pathology following acute JHMV-induced encephalitis

To first determine the consequences of plaque deposition on cell types within the CNS, we performed DGE analysis on control 5xFAD and control WT brains. Volcano plots demonstrate the widespread transcriptional changes within each CNS subtype and major cell type impacted by plaque deposition **(Supplementary Figure 7 & 8A)**. In calculating the DEG score to compare the magnitude of fold change in DEGs within different CNS cell types, we found that myeloid cells demonstrated the greatest magnitude of down- and up-regulated DEGs compared to other CNS cell types. Displaying the DEG score of each cell in XY space further depicts the distribution of dysregulated gene expression across myeloid cells in plaque-laden areas, particularly the cortex and subiculum of 5xFAD mice **(Supplementary Figure 8B&C)**. In total, the presence of amyloid plaque deposition resulted in 190 significantly down-regulated DEGs within myeloid cells (defined as p_adj_ < 0.05 and absolute average difference > 0.3) between 5xFAD and WT brains. In line with previous sequencing studies, myeloid cells impacted by the presence of amyloid plaque pathology exhibited down-regulated homeostatic genes (*P2ry12, Tmem119, Csf1r)*. Additionally, pathway analysis via gene ontology revealed down-regulation in pathways involved in modulation of neurotransmission, cell morphogenesis, cell migration, development of neuron projections, and learning and memory **(Supplementary Figure 8D)**. Furthermore, myeloid cells exhibited 77 significantly up-regulated DEGs between 5xFAD and WT brains, several of which are associated with disease-associated microglia (DAM) signatures (*Cst7, Itgax, Spp1, Gpnmb, Ctss, Ctsz, Csf1, Lpl, etc.)* **(Supplementary Figure 8A)**. Meanwhile, additional up-regulated DEGs within myeloid cells were associated with GO pathways involving regulation of tumor necrosis factor and interleukin-6 production, gliogenesis and glial cell proliferation, and negative regulation of cell activation **(Supplementary Figure 8E).**

We next performed differential gene expression (DGE) analysis to investigate how the acute viral encephalitis impacted CNS cell types surrounding Aβ plaque pathology. Non-aggregate DGE analysis of JHMV-infected 5xFAD vs. 5xFAD brains revealed dysregulated gene expression across the 42 cell clusters and major cell types **(Supplementary Figure 9 & Figure 5E)**. To visualize the spatial distribution of all cell types with changes in gene expression, we generated a DEG score representing the magnitude of fold change between each comparison of two experimental groups **(Figures 5F&G)**. Visualizing down-regulated DEG scores across brain sections demonstrates widespread genetic changes during acute JHMV-induced encephalitis, however inhibitory and excitatory neuron clusters exhibited the greatest DD score. Replotting the inhibitory neuron clusters in XY space validated the correct cell type clustering with spatial specificity. To visualize genetic changes in inhibitory neurons across the different experimental groups, we plotted the DEGs within this cluster as pseudo-bulked expression values and in XY space **(Figure 5H&I)**. Within inhibitory neurons in JHMV-infected 5XFAD brains, several genes were downregulated involved in memory, cognition, and behavior such as *Cck, Cnr1, Vip, Sgk1,* and *Egr1* **(Figure 5J)**.

Among all the cell types, myeloid cells expressed the greatest differential upregulation (DU) score between JHMV-infected 5xFAD mice and uninfected 5xFAD controls, implicating a post-acute encephalitis response that may interact with existing amyloid pathology. We observed an increased proportion of myeloid cells during viral encephalitis in 5xFAD brains, in concordance with increased infiltration following JHMV infection^28^ **(Figure 5D)**. Using DGE analysis, we found that several genes associated with immune activation and antigen presentation are upregulated in the myeloid cell cluster (*CD74, H2-Aa, H2-Ab1;* **Figure 5E**). Interestingly, several DAM genes were downregulated within this myeloid population (*CD9, Trem2, Clec7a, Ctsz, Dtsd;* **Figure 5E**), as well as several homeostatic microglial genes (*Csf1r, P2ry12, Hexb, Cst3;* **Figure 5E**). To determine the biological pathways enriched in myeloid DEGs, we performed gene ontology (GO) pathway analysis of dysregulated genes in the myeloid cluster. We found that JHMV-infection induced several up-regulated myeloid DEGs in 5xFAD brains associated with gliogenesis, glial cell development, glial cell differentiation, inflammatory responses, and myelination **(Supplementary Figure 10A)**. We also utilized protein network analysis to illustrate the protein-protein interactions of closely associated up-regulated myeloid DEGs **(Supplementary Figure 10B)**. Interestingly, GO pathway analysis of down-regulated myeloid DEGs depicted enriched pathways involved in regulation of macrophage migration, host-pathogen processes, ischemia responses, and neuronal death **(Supplementary Figure 10C&D)**.

Pseudo-bulk analysis of these top DEGs between JHMV-infected 5xFAD and 5xFAD brains was performed, demonstrating the downregulation of several DAM genes following acute viral encephalitis in 5xFAD myeloid cells despite DAM upregulation relative to control WT myeloid cells **(Figure 6A)**. Cell clustering of all sampled cells produced four subtypes within the myeloid cluster: DAM, Activated, Myeloid 1, and Myeloid 2. Plotting these clusters in XY space reveals that the DAM and Myeloid 1 subtype to be distributed in plaque-laden areas (i.e. the subiculum of the hippocampus and throughout the cortex; **Figure 6B, Supplementary Figures 11A&D)**. Meanwhile, the Activated Myeloid cluster appears to be predominantly distributed throughout the brain in regions associated with viral replication, whereas the Myeloid 2 population is primarily localized to white-matter (i.e. the corpus callosum; **Figure 6B, Supplementary Figures 11B&C)**. Concordant with our histology and flow cytometry analysis, we found an increased proportion of cells within the DAM and Activated Myeloid subtype following acute viral encephalitis in both 5xFAD and WT brains **(Figure 6C)**.

**FIGURE 6.**
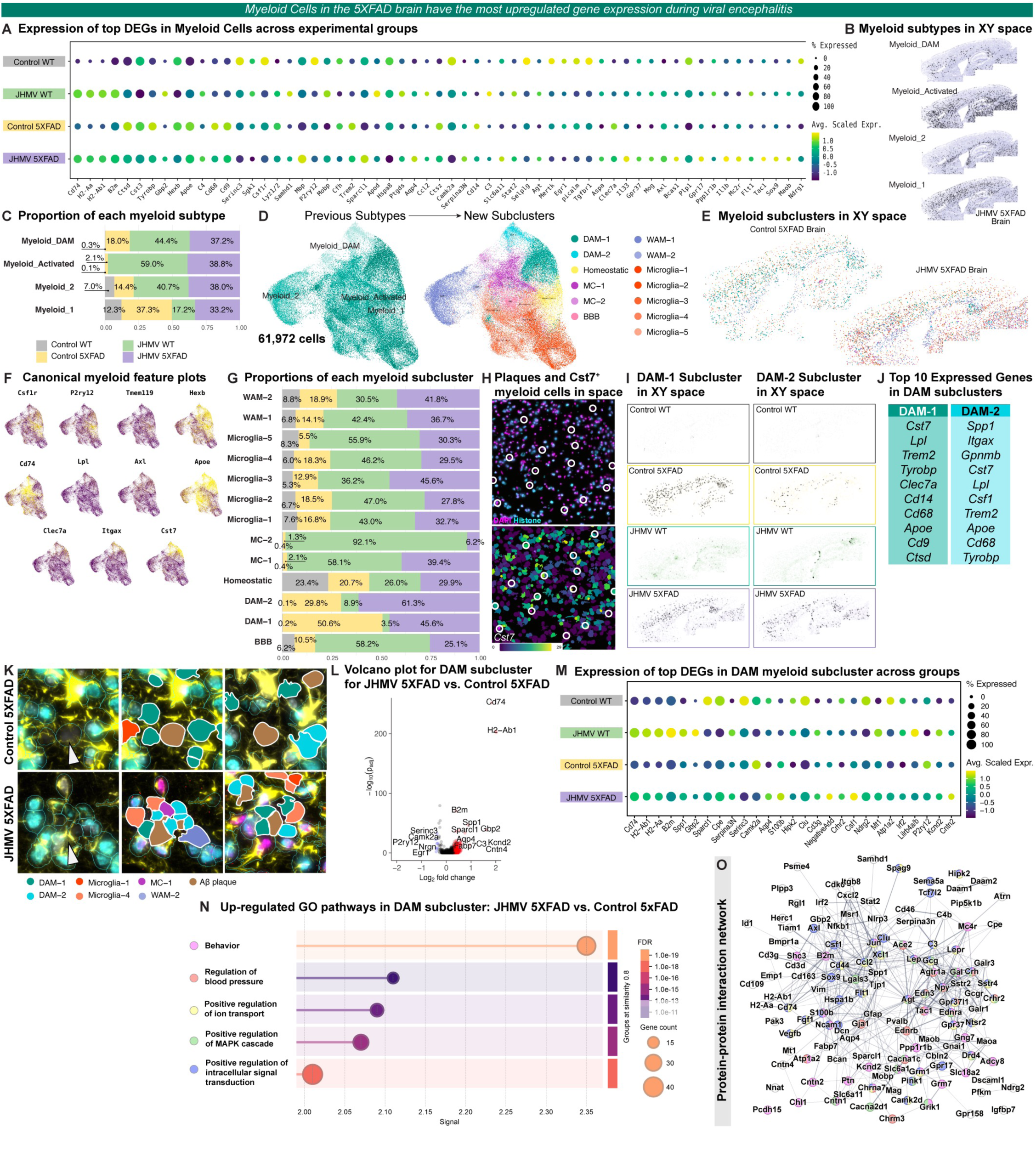
Sub-clustering analysis of myeloid cells. **A)** Expression of top DEGs in myeloid cells between JHMV-infected 5xFAD vs. control 5xFAD across each experimental group. **B)** Myeloid cell subtypes in XY space on a representative section of a JHMV-infected 5xFAD brain. **C)** Proportion of each myeloid cell subtype split by each experimental group. **D)** Myeloid cells were re-clustered at 0.5 resolution yield 13 new myeloid subclusters. UMAP of 61,972 subsetted myeloid cells with original myeloid annotations (left) and new myeloid subcluster annotations (right). **E)** Myeloid cell subclusters in XY space on a representative control 5xFAD and JHMV-infected 5xFAD brain section. **F)** Feature plots representing expression of several canonical myeloid cell markers on the new UMAP. **G)** Proportion of the number of cells in each myeloid subcluster, grouped by experimental group. **H)** Representative FOV of DAPI^+^ plaques and Histone-expressing cells in the subiculum region of JHMV-infected 5xFAD mouse (top). Cst7-expressing cells in the same FOV (below). **I)** DAM-1 (left) and DAM-2 (right) subclusters in XY space on representative sections from each experimental group. **J)** List of the top 10 expressed genes in DAM-1 (left) or DAM-2 (right) subclusters. **K)** Representative FOVs of subiculum in JHMV-infected 5xFAD (top) and control 5xFAD (bottom) with arrows pointing to DAPI-positive amyloid-beta dense core plaques surrounded by myeloid subtypes. **L)** Volcano plot for DAM subcluster in JHMV-infected 5xFAD vs. control 5xFAD. **M)** Pseudo-bulk sequencing analysis of top DEGs for JHMV-infected 5xFAD and control 5xFAD within DAM myeloid subcluster. **N)** Pathway analysis by Gene Ontology of enriched DEGs in the DAM myeloid subcluster for JHMV-infected 5xFAD vs. control 5xFAD comparison. **O)** Protein-protein interactions (PPI) network of all significantly up-regulated DEGs within DAM myeloid subcluster in JHMV-infected 5xFAD vs control 5xFAD. Nodes are colored according to enriched pathway analysis. The thickness of each line connection between nodes indicates the degree of confidence in the prediction of the PPI.

To further investigate the myeloid population, we captured all 61,972 myeloid cells across the four experimental groups and re-clustered them into 13 new subclusters **(Figure 6D)**. We performed manual cell annotation once more using top gene expression and their spatial location in the brain **(Figure 6E, Supplementary Figure 11E)**. Among the sub-clustered myeloid cells, we identified two subclusters within the DAM subtype (DAM-1 and DAM-2). Visualization of cellular expression of canonical DAM markers *Lpl, Axl, Clec7a, Itgax, and Cst7* corroborates the identification of these DAM subclusters **(Figure 6F)**. Cell proportions were calculated for each myeloid sub-cluster across the four experimental groups, revealing a higher proportion of both DAM sub-clusters are represented in 5xFAD brains **(Figure 6G, Supplementary Figure 11F-I)**. While the DAM-1 sub-cluster is equally represented in both JHMV-infected and control 5xFAD brains, we observed a greater proportion of cells in the DAM-2 sub-cluster in 5xFAD brains during viral encephalitis **(Figure 6G, Supplementary Figure 9I)**. In the cell segmentation imaging, we identified several DAPI-positive “cells” that were only present in 5xFAD brains, anucleate, and lacked expression of histone markers, suggesting these to be DAPI-positive Aβ plaques surrounded by *Cst7*-expressing myeloid cells **(Figure 6H)**.

Plotting these DAM subclusters in XY space demonstrates that both DAM-1 and DAM-2 subclusters are distributed throughout the 5xFAD brain in plaque-laden areas, representing plaque-associated myeloid cells **(Figure 6I)**. Furthermore, both DAM-1 and DAM-2 sub-clusters are characterized by high expression of canonical DAM markers (*Cst7, Lpl, Trem2, Apoe)* and phagocytosis (*Tyrobp, CD68),* likely attributed to plaque-associated microglia. However, cells within the DAM-1 sub-cluster differs in its high expression of *Clec7a, Cd9, and Cd14* while the DAM-2 sub-cluster exhibit a high expression of *Itgax, Csf1, Spp1, and Gpnmb* **(Figure 6J)**. To determine the association amyloid pathology and cells with each DAM sub-cluster, we examined the proximity of DAM-1 and DAM-2 cells surrounding Aβ plaques that stained positive DAPI but lacked histone markers **(Figure 6K)**. We observed several DAM-1 cells surrounding Aβ plaques in control 5xFAD mice, corroborating that DAM-1 likely represent plaque-associated microglia. Meanwhile, we confirmed a greater proportion of DAM-2 cells surrounding Aβ plaques in JHMV-infected 5xFAD brains. Furthermore, these DAM-2 cells appear to be in direct contact with dense-core Aβ plaque pathology following encephalitis, unlike DAM-1 cells in control 5xFAD brains **(Figure 6K)**. This data appears in line with our histological data, suggesting that cells within the DAM-2 sub-cluster could be responsible for the higher association of inflammatory MAC2^+^ myeloid cells surrounding Aβ plaques in response to JHMV infection.

To investigate the transcriptional changes that may occur in cells within the DAM sub-cluster, we performed pseudo-bulk analysis of DEGs across all the experimental groups **(Figures 6L&M, Supplementary Figure 12A)**. Relative to control 5xFAD brains, cells in the DAM sub-cluster had increased expression of genes associated with immune activation and antigen-presentation *(CD74, H2-Ab1, H2-Aa, B2m)* and IFN-γ-induced signaling *(Gbp2, Irf2)* **(Figures 6L&M)**. Gene ontology pathway analysis of these up-regulated DEGs in DAMs depict effects on behavior, regulation of blood pressure, ion transport, regulation of MAPK cascade, and regulation of intracellular signal transduction **(Figure 6N)**. To visualize the protein-protein interactions and pathways associated with significant DEGs in DAM cells following viral encephalitis, we visualized interacting networks of related proteins expressed by up-regulated DEGs **(Figure 6O)**. Furthermore, performing DGE analysis on each DAM sub-cluster reveals a differential impact of JHMV infection on DAM-1 and DAM-2 sub-clusters in the 5xFAD brain **(Supplementary Figures 12B&C)**. In the DAM-1 subcluster, pathway analysis illustrates enriched pathways for up-regulated DEGs involved in regulation of MAPK cascade, gliogenesis, blood pressure, and secretion **(Supplementary Figure 12D)**. Down-regulated genes within DAM-2 cells enrich GO pathways involved in regulation of macrophage fusion, microglial cell activation, and responses to lipoprotein particles, while up-regulated genes are associated with antigen presentation via MHC Class II, T cell activation, and regulation of adaptive immunity **(Supplementary Figure 12E)**.

Corroborating previous studies detailing the infiltration of monocyte-derived macrophages during JHMV-induced encephalitis^28^, we have also observed high proportions of monocyte-derived cells (MC) within JHMV-infected WT and JHMV-infected 5xFAD brains in our spatial transcriptomic myeloid dataset **(Figure 6G, Supplementary Figure 12F)**. Plotting the MC subclusters in XY space illustrates a significant spread of MC-1 throughout the brains of JHMV-infected mice, yet MC-2 appears more specific to JHMV-infected WT mice **(Supplementary Figure 12G)**. When comparing transcriptional differences of monocyte-derived cells within the MC subcluster between JHMV-infected brains vs. Control 5xFAD brains, we find that JHMV infection results in decreased expression of *Lyz1, Mrc1, Tyrobp, Clec7a, Cd44,* and *Ctsd* **(Supplementary Figure 12H&I).** Utilizing pseudo-bulk sequencing analysis, we have also identified dysregulated DEGs among other myeloid cell types: WAM, Homeostatic myeloid cells, BBB-associated myeloid cells, and microglia **(Supplementary Figure 13)**.

### Comparison of myeloid differential gene expression effects of Aβ pathology and JHMV-induced encephalitis

We have demonstrated that acute viral encephalitis through JHMV infection shifts the transcriptional signature, particularly in disease-associated myeloid cells surrounding Aβ plaques. To investigate whether dysregulated responses in myeloid cells is a consequence of encephalitis, the presence of Aβ pathology, or unique to the presence of both inflammatory stimuli, we compared the average differences in gene expression within all myeloid cells in our spatial transcriptomic dataset. We plotted all DEGs between Control 5xFAD vs. Control WT (response to amyloid pathology) and JHMV WT vs. Control WT (response to viral encephalitis) using the average difference of each gene among the two comparisons **(Figure 7A)**. Many gene expression changes were shared and correlated between the two inflammatory stimuli. In response to either amyloid or JHMV encephalitis, myeloid cells shared down-regulation of 59 genes with similar homeostatic markers (e.g., *P2ry12, Cx3cr1, Tmem119, Csf1r, etc.*), while sharing up-regulation in 47 genes related to DAM signatures (*e.g., Apoe, Clec7a, Cst7, Ctsb, Itgax, Axl, etc.).* Notably, several genes were more upregulated in response to one stimulus such as *H2-Aa, Cd74, H2-Ab1, Gpnmb, Lgals3*, which demonstrated greater upregulation in response to JHMV infection **(Figure 7B)**. Meanwhile, there were several genes that were uniquely differentially expressed in one inflammatory stimulus over the other. For example, *Cd9, Trem2, Cd68, C1qb, C1qc* were upregulated within myeloid cells in response to amyloid pathology compared to JHMV infection. Conversely, *Cd14, Cst3, Picalm, Abi3, Bin1, Kif5c* were more downregulated in response to viral encephalitis compared to amyloid pathology **(Figure 7B)**. Therefore, while the activation response to viral encephalitis and amyloid pathology similarly upregulated DAM-related, inflammation genes, there were nuanced differences in related groups of genes when considering the magnitude of their inflammatory response.

**FIGURE 7.**
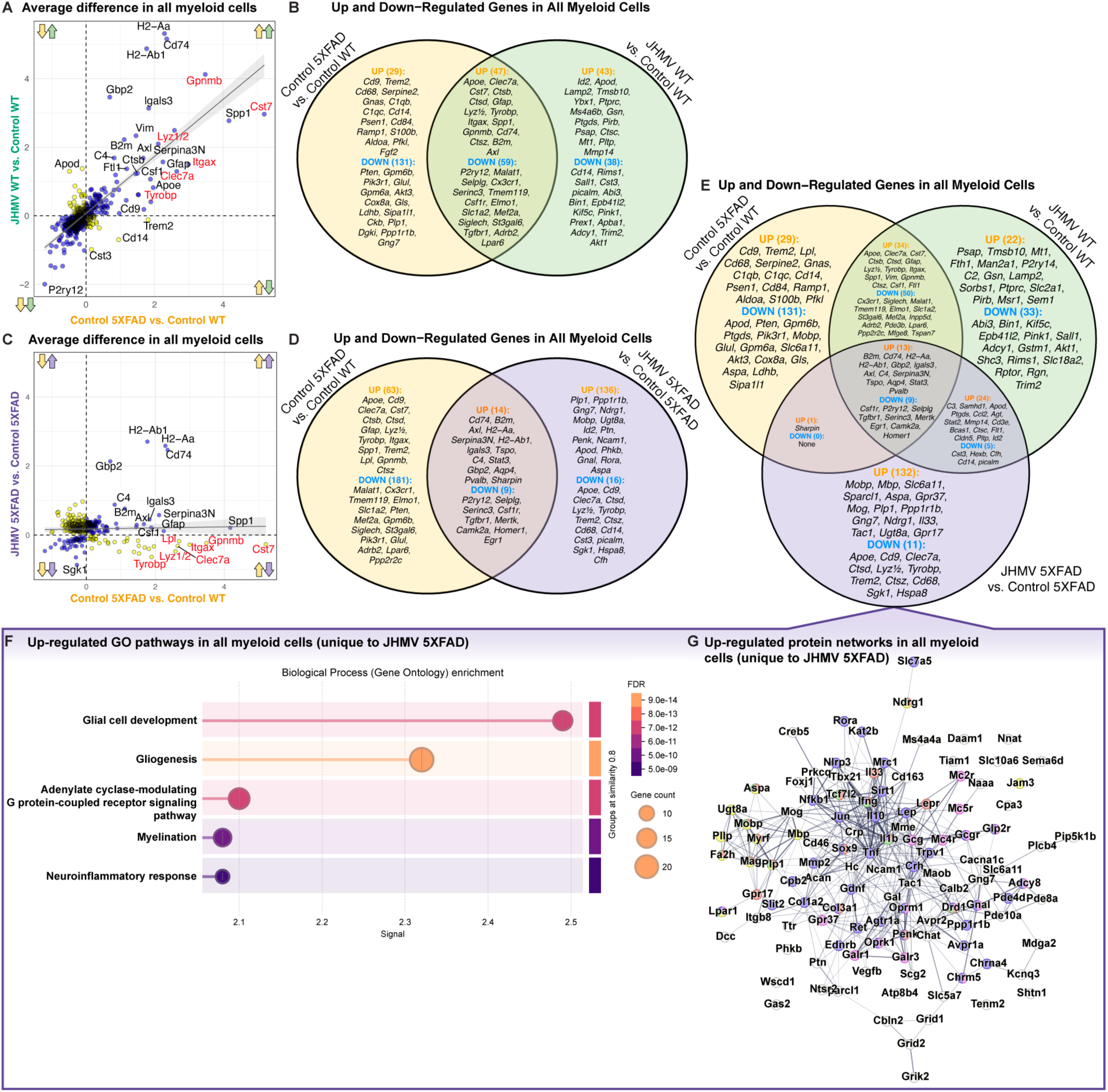
Myeloid cells have differential responses to the presence of viral encephalitis, amyloid pathology, or both concurrently. **A)** Scatterplot of the average difference of all myeloid cells for all significant genes (padj < 0.05) between Control 5xFAD vs. Control WT (x-axis, only amyloid pathology) and JHMV WT vs. Control WT (y-axis, only viral encephalitis) comparisons. Arrows indicate direction of dysregulation for each comparison. Directly correlated genes (blue) occur in the same direction for both comparisons (i.e., both up-regulated or both down-regulated), while inversely correlated genes (orange) occur in opposite directions for each comparison. Linear regression line demonstrates the relationship between the two comparisons. **B)** Venn diagram depicting the significant up- and down-regulated genes unique to the Control 5xFAD vs. Control WT (left, yellow) and JHMV WT vs. Control WT (right, green), while demonstrating up- and down-regulated genes commonly shared across the two comparisons (middle). **C)** Scatterplot of the average difference of all myeloid cells for all significant genes between Control 5xFAD vs. Control WT, now with JHMV 5xFAD vs. Control 5xFAD (y-axis, both amyloid and viral encephalitis). Red text demonstrates genes up-regulated in the same direction in the first scatterplot, but down-regulated in the JHMV 5xFAD vs. Control 5xFAD comparison. **D)** Venn diagram depicting the significant up- and down-regulated genes unique to Control 5xFAD vs. Control WT and JHMV 5xFAD vs. Control 5xFAD (right, purple), while demonstrating up- and down-regulated genes commonly shared between the two comparisons. **E)** Three-way Venn diagram of all significantly up- and down-regulated genes between the three comparisons: Control 5xFAD vs. Control WT (yellow), JHMV WT vs. Control WT (green), and JHMV 5xFAD vs. Control 5xFAD (purple). Genes are considered significantly correlated if the log2 Fold Change magnitude is greater than 0.3. **F)** Gene ontology pathways enriched by all significantly up-regulated DEGs unique to just the JHMV 5xFAD vs. Control 5xFAD comparison. **G)** PPI of the significantly up-regulated DEGs unique to the JHMV 5xFAD vs. Control 5xFAD comparison. Nodes are colored according to enriched pathway analysis. The thickness of each line connection between nodes indicates the degree of confidence in the prediction of the PPI.

To determine the effect of the concurrence of both inflammatory stimuli (e.g., amyloid pathology and viral encephalitis) on the gene expression within myeloid cells, we plotted all DEGs between Control 5xFAD vs. Control WT and JHMV 5xFAD vs. Control 5xFAD **(Figure 7C)**. Once again, many differential gene expression changes were shared in the context of only amyloid pathology (Control 5xFAD) or *both* amyloid and viral encephalitis. Myeloid cells shared 14 up-regulated genes involving MHC class II presentation, adaptive immunity activation, and other inflammatory mechanisms like *Cd74, B2m, H2-Aa, H2-Ab1, Stat3, Gbp2* **(Figure 7D)**. Additionally, 9 homeostatic microglia genes were similarly downregulated in both conditions, such as *P2ry12 and Csf1r*.

Notably, to determine which set of DEGs were unique to the interaction of both amyloid pathology and viral encephalitis, we compared DEGs from all myeloid cells across three comparisons: Control 5XFAD vs. Control WT (only amyloid pathology), JHMV WT vs. Control WT (only JHMV infection), or JHMV 5XFAD vs. Control 5XFAD (both amyloid pathology and JHMV infection). We have identified 143 genes that uniquely become differentially expressed under the occurrence of both amyloid pathology and JHMV infection. Several DAM genes are among the 11 uniquely downregulated in the interaction of both inflammatory stimuli in JHMV-infected 5xFAD brains: *Apoe, Cd9, Clec7a, Ctsd, Lyz1/2, Tyrobp, Trem2, Ctsz, and Cd68*. Notably, these same set of DAM genes were observed to be upregulated in the presence of both amyloid pathology and viral encephalitis, independently **(Figure 7F)**. Additionally, 132 genes were observed to be uniquely upregulated in the interaction of both inflammatory stimuli **(Figure 7F)**. Pathway analysis of these up-regulated genes in JHMV 5XFAD brains reveal impacts on glial cell development, gliogenesis, adenylate cyclase-modulating G protein-coupled receptor signaling pathways, myelination, and neuroinflammatory responses **(Figure 7G)**. Additionally, mapping out protein-protein interactions of significantly up-regulated DEGs unique to JHMV 5xFAD myeloid cells demonstrate closely linked genes that transcribe proteins associated with interactions or similar cellular mechanisms.

## Discussion

With increasing evidence identifying how Aβ-dependent pathways in combination with innate immune responses influence AD onset in pre-clinical mouse models of AD, microbial infection and its resulting peripheral inflammation has been put forward as a major environmental risk factor for AD onset^11,12,35–38^. Importantly, viral exposure has recently been associated with increased risk of numerous neurodegenerative diseases, with the strongest association occurring between viral encephalitis and AD^10^. Viral encephalitis is defined as viral-mediated inflammation of the brain parenchyma. While the BBB prevents most viral pathogens from invading from the periphery, neurotropic viruses can circumvent these barriers to infect and replicate within the CNS while the effects of neurovirulent viruses replicating in the periphery can indirectly induce neurological effects^11^. The resulting encephalitis can therefore lead to neurologic damage and potential consequences to the onset or severity of neurodegenerative disease^12–14^. There is a critical need to understand post-infectious neuroimmune mechanisms and how they may interact with Aβ plaque deposition and modulate the onset and severity of AD pathology.

Using the neurotropic JHM strain of mouse hepatitis virus (JHMV, a *Betacoronavirus* genus member, we demonstrate that acute JHMV infection in 5xFAD mice results in similar inflammatory responses in parallel to significant reductions in Aβ plaque sizes despite no significant changes to overall Aβ plaque load. We also observed localized interactions between infiltrating monocyte/macrophages and dense-core Aβ plaques in JHMV-infected 5xFAD brains. Within regions where viral RNA/antigen persists, increased recruitment of inflammatory monocyte/macrophages and MAC2^+^ macrophages significantly correlate with more compact Aβ plaques. Furthermore, we utilized spatial transcriptomic imaging to investigate the transcriptional impact of viral encephalitis with single-cell resolution and determine altered gene expression within specific cell types surrounding Aβ pathology in the 5xFAD brain. While several cell types were dysregulated following viral encephalitis, we show that myeloid cells demonstrate a suppressed DAM response in the brain of JHMV-infected 5xFAD mice, particularly those surrounding amyloid pathology.

During neurodegenerative diseases like AD, microglia transition from homeostatic states into activated inflammatory states, which correlate with cognitive decline and neurodegeneration^39,40^. Single-cell transcriptomic and proteomic studies have revealed activated microglia further assume functionally distinct microglial cell populations in during different diseased states within the CNS. In AD, the prominent transcriptional signature within microglia are disease-associated microglia (DAMs), microglia neurodegenerative phenotype (MGnD), and activated response microglia (ARMs)^8,40^. The functional role of many upregulated DAM genes are also implicated in phagocytosis, chemotaxis, damage-mediated cytokine release, while downregulated genes are typical of microglial homeostasis^40^. During early, asymptomatic stages, increased microglia activation into DAM states can promote Aβ clearance and confer neuroprotection, which can shift to a neurodegenerative, proinflammatory state reminiscent of the MGnD phenotype as disease progresses further^41^.

Other myeloid cells contributing to neuroinflammation are non-parenchymal macrophages, which undergo development from bone marrow (BM)-derived monocytes and represent functionally distinct cell populations from microglia under steady-state conditions^42–44^. However, studies have reported that inflammatory and diseased conditions (e.g., AD) result in robust infiltration of peripheral-derived monocytes into the brain, which differentiate into macrophages upon immune activation with similar transcriptional and phenotype changes as microglia^34,45–48^. In post-mortem AD tissue, circulating monocyte/macrophages have been identified, infiltrating into brain parenchyma and localizing amyloid plaques^47^. In pre-clinical AD models, infiltration of peripheral BM-derived monocyte/macrophages have also been found closely associating with Aβ plaques^34,49,50^. Notably, these BM-derived macrophages appear to be more efficient than resident microglia in phagocytosis and clearance of Aβ within the brain parenchyma, while protecting synapses from loss^50–52^. The ability of BM-derived macrophages to reduce Aβ is further illustrated in selective ablation studies resulting in exacerbated cortical and hippocampal Aβ pathology. Conversely, disease stage may impact the phagocytic capacity of peripheral myeloid cells in clearing Aβ during the late-stage disease^53^. Further supporting the importance of timing monocyte and macrophage infiltration, glatiramer acetate used to recruit immunomodulatory BMDMs only modestly reduced Aβ burden in 5xFAD mice during the development stage of pathology and increased Aβ levels during later stages of neuropathology^54^. These results provide additional evidence of the importance of myeloid cells in regulating neuroinflammation and shaping AD pathology, specifically highlighting the nuanced disease-modifying role of peripheral MDMs recruited into the CNS. These results provide additional evidence of the importance of myeloid cells like monocyte/macrophages in regulating neuroinflammation and shaping AD pathology^55,56^.

This study builds upon an ongoing question in the field about the role of infectious microbes, its result inflammation, and its effects on the progression and severity of AD pathologies. Previously, our lab has reported on the impact of JHMV encephalitis on the 3xTg-AD transgenic mouse model, which uniquely combines the autosomal dominant Swedish mutation in human *APP* to reproduce amyloid pathology and two human *MAPT* mutations associated with frontotemporal dementia (FTD) to reproduce tauopathies^57^. While JHMV infection did not impact Aβ pathology in 3xTg-AD brains, the resulting encephalitis did appear to worsen tau pathology^57^. Other viral models have also been explored as potential risk factors for AD. Post-mortem analysis has identified colocalization of HSV-1 viral DNA with Aβ plaques in AD brains^58^. This is also supported with a study reporting that HSV-1 infection led to seeding of surface glycoproteins within Aβ oligomers and accelerated Aβ deposition in the 5xFAD mouse model of familial AD, suggesting Aβ playing a potentially protective role in anti-viral immunity^59^. However, recent studies have also complicated this prevailing theory in which unique post-mortem AD brain tissue with concurrent HSV encephalitis revealed HSV-infected cells localized close to Aβ plaques neurofibrillary tangles, but not directly associated with worsening AD pathology^60^. Additionally, 5xFAD nor combined LOAD mutations (e.g., humanized Aβ, *APOE4*, and *TREM2*^R47H^ transgenes) in mice did not significantly protect against HSV-1 exposure nor initiate Aβ aggregation following HSV-1 infection, respectively^61,62^. Interestingly, the researchers did observe limited viral invasion in regions with high Aβ plaque densities attributed with phagocytic activity of reactive microglia^61^. Additionally, HSV-1 infection of LOAD mice resulted in extensive infiltration of peripheral leukocytes and phagocytic activity of myeloid cells^62^. Rather than a direct cause, HSV-1 infection may be a modulating risk factor for AD with a combination of both environmental and genetic factors contributing to neurological damage.

While this study reports a dampened DAM transcriptional response to amyloid pathologies following encephalitis, there are several limitations that warrant future investigations. Here, we utilize single-cell spatial transcriptomic imaging as a powerful method of retaining both spatial and transcriptional information within unique cell types. However, the pre-selected 1000-plex mouse neuroscience probe list limits the depth of exploration and unbiased investigation compared to whole genome approaches offered by traditional single-cell and single-nucleus techniques. Specifically, the available 1000-plex probe list offers probes against key gene transcripts involved in major biologically-relevant pathways, yet the limited selection of probes for inflammation and immunity pathways yield surface-level analyses on immune cell types. Interestingly, DGE analysis of each CNS cell type in the spatial transcriptomic dataset illustrates up-regulation of antigen presentation genes *H2-Aa* and *H2-Ab1*, even among cells that are not typically associated with antigen presentation. These are likely due to segmentation overlaps with antigen-presenting cells such as microglia and CNS-associated macrophages, highlighting some limitations in the cell segmentation process for identifying captured cells within the tissue sample. Furthermore, it will be necessary to carefully consider and interpret the impact on pathologies observed in the 5xFAD transgenic model, which lacks AD-related tauopathies. While we have previously investigated the impact of viral encephalitis on a mouse model with both amyloidosis and tauopathy, its reliance on transgenes to overexpress rare autosomal-dominant mutations limits the clinical relevance of this approach. With the advent of better humanized mouse models that have the potential of recapitulating amyloidosis and tauopathy together, future investigations investigate both the acute and post-acute impact of viral encephalitis on these mice may offer more relevant and comprehensive understanding of its interactions with AD pathologies and its severity. Committing to defining these post-infectious neuroimmune mechanisms and their interaction with AD pathogenesis holds promise, with potential to offer both treatments for current patients and preventative interventions to improve public health outcomes and reducing AD risk in an aging population.

## Concluding Remarks

Ongoing studies have demonstrated a relationship between viral encephalitis and Alzheimer’s disease. This study was performed to better elucidate the interaction between viral-induced inflammation and amyloid pathology associated with Alzheimer’s disease using experimental infection of aged 5xFAD and WT mice with JHMV, a neurotropic murine coronavirus. We found that JHMV infection induced infiltration of MAC2^+^ macrophages and T cells in the brains of 5xFAD mice. While viral encephalitis did not significantly alter overall Aβ burden, we observed reductions in dense-core Aβ plaque size and numbers associated with MAC2^+^ macrophages within subiculum and somatosensory cortex regions, respectively. Therefore, we employed spatial transcriptomic imaging to parse out transcriptional changes within neighboring cells surrounding Aβ plaques with single cell resolution. In doing so, we found significant transcriptional dysregulation within myeloid cells in JHMV-infected 5xFAD brains, specifically impacting genes associated with DAM responses to plaque pathology. These findings reveal that viral encephalitis and its resulting inflammatory mechanisms impacts the expression of key genes within myeloid cells responding to Aβ plaque pathology, which has implications to the role of viral encephalitis as a risk factor for Alzheimer’s disease.

## Funding Information

Alzheimer’s Association grant 105190 (TEL)

National Institutes of Health (NIH-NINDS) grant R35NS116835 (TEL)

National Institutes of Health (NIH-NIA) grant R01AG081599 (KNG)

National Institutes of Health (NIH-NIA) grant U54 AG054349 (KNG)

National Institutes of Health (NIH-NINDS) Training Grant T32 NS121727 (DIJ, NK)

National Institutes of Health (NIH-NINDS) F31NS141599 (KIT)

## Author Contribution Statement

D.I.J., K.N.G., and T.E.L. designed the study; S.F., L.L., K.F., J.M., V.S., and R.J. helped D.I.J. perform experiments; D.I.J. analyzed experimental data; D.I.J. performed bulk RNA sequencing analysis; D.I.J. performed spatial transcriptomics; D.I.J. analyzed spatial transcriptomic data; K.I.T. and N.E.K. provided technical advice and scripts for spatial transcriptomic analysis.

## Data Availability Statement

Protocols, data and results will be available via the AD Knowledge Portal.

