## Supplementary Table for "Characterization of effects of a neurotropic murine coronavirus infection on Alzheimer’s disease neuropathology of 5xFAD mice"

| **Primary Antibody** | **Source** | **Catalog Number** | **Dilution** | **Secondary Antibody** | **Source** | **Catalog Number** | **Dilution** |
| --- | --- | --- | --- | --- | --- | --- | --- |
| IBA1 | Wako | 09-19741 | 1:2000 | Goat anti-rabbit IgG, Alexa Fluor 488 | ThermoFisher | A11008 | 1:200 |
| MAC2 | Cedarlane | CL8942AP | 1:500 | Goat anti-rat IgG Alexa Fluor 594 | ThermoFisher | ab150160 | 1:200 |
| 6e10 Aβ1-16 | BioLegend | 8030001 | 1:2000 | Goat anti-mouse, Alexa Fluor 555 | ThermoFisher | A21424 | 1:200 |
| OC | Sigma Aldrich | 2286 | 1:1000 | Goat anti-rabbit IgG, Alexa Fluor 488 | ThermoFisher | A11008 | 1:200 |
| CD4 | Abcam | ab183685 | 1:500 | Goat anti-rabbit IgG, Alexa Fluor 488 | ThermoFisher | A11008 | 1:200 |
| CD8 | Abcam | ab217344 | 1:500 | Goat anti-rabbit IgG, Alexa Fluor 488 | ThermoFisher | A11008 | 1:200 |
