## Supplementary Figures for "Characterization of effects of a neurotropic murine coronavirus infection on Alzheimer’s disease neuropathology of 5xFAD mice"

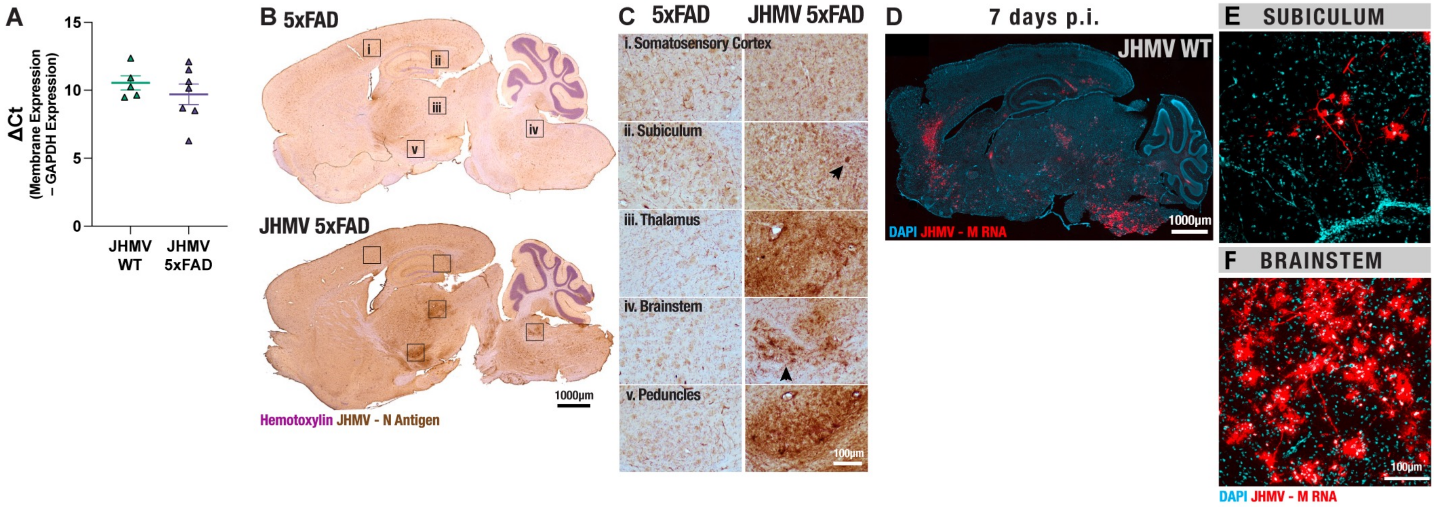
A
ΔCt
(Membrane Expression – GAPDH Expression)
15
10
5
0
JHMV WT
JHMV 5xFAD
B 5xFAD
i
ii
iii
iv
v
JHMV 5xFAD
1000μm
Hemotoxylin JHMV - N Antigen
C 5xFAD
JHMV 5xFAD
i. Somatosensory Cortex
ii. Subiculum
iii. Thalamus
iv. Brainstem
v. Peduncles
100μm
D
7 days p.i.
JHMV WT
1000μm
DAPI JHMV - M RNA
E SUBICULUM
F BRAINSTEM
100μm
DAPI JHMV - M RNA

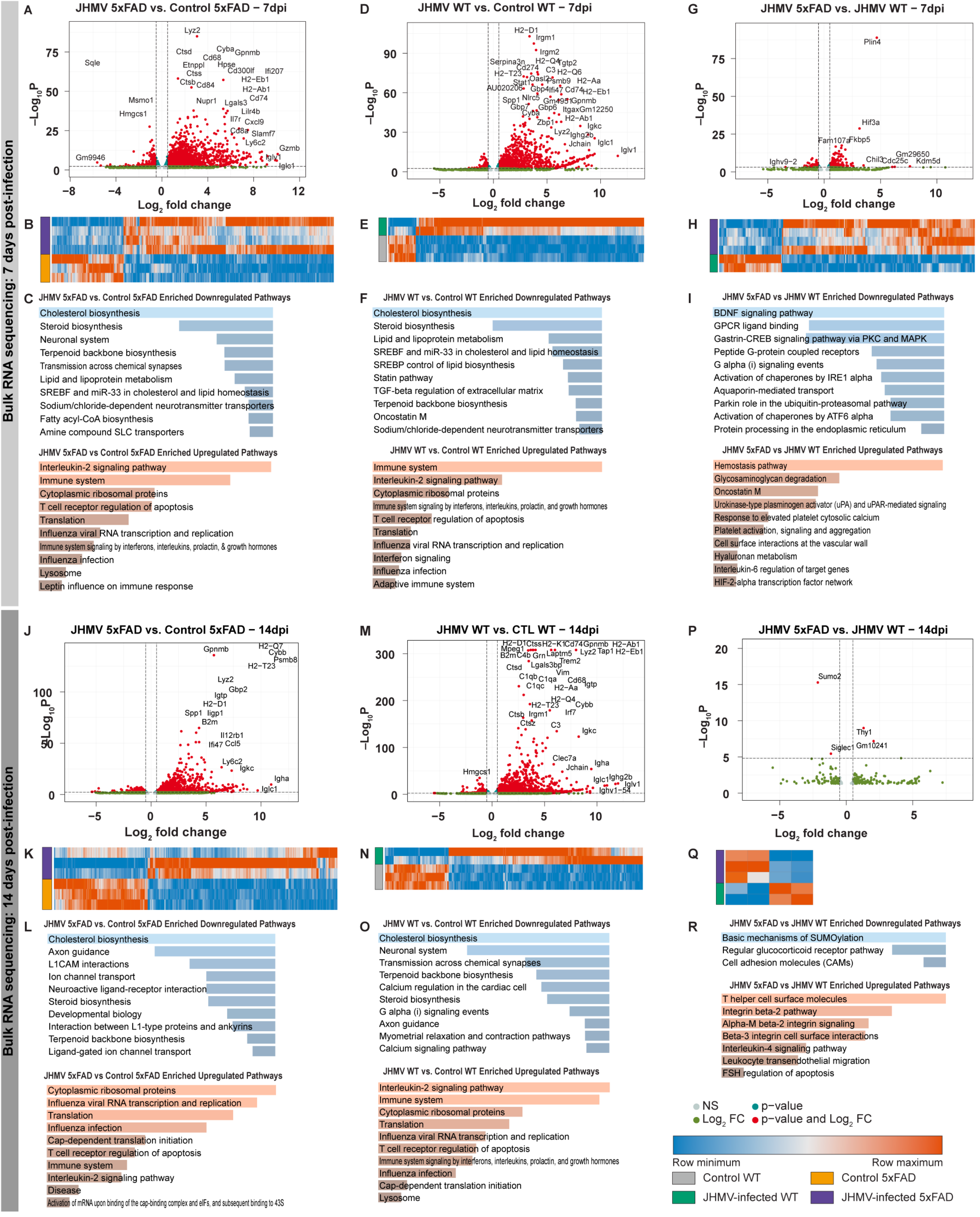
Bulk RNA sequencing: 7 days post-infection
A
JHMV 5xFAD vs. Control 5xFAD – 7dpi
Lyz2
Ctsd
Cyba
Gpnmb
Cd68
Sqle
Etnppl
Hpse
Ctss
Cd300lf
Ifi207
Ctsb
Cd84
H2−Eb1
H2−Ab1
Msmo1
Nupr1
Lgals3
Cd74
Hmgcs1
Lilr4b
Il7r
Cxcl9
Cd8a
Slamf7
Ly6c2
Gzmb
Gm9946
Iglv1
Iglc1
−Log10 P
Log2 fold change
D
JHMV WT vs. Control WT – 7dpi
H2−D1
Irgm1
Irgm2
Serpina3n
H2−Q4
Tgtp2
Cd274
H2−T23
C3
H2−Q6
Oasl2
Psmb9
H2−Aa
Stat1
AU020206
Gbp4
Ifi47
Cd74
H2−Eb1
Spp1
Nlrc5
Gm4951
Gpnmb
Gbp7
Gbp6
Cyba
Itgax
Gm12250
H2−Ab1
Zbp1
Igkc
Lyz2
Ighg2b
Jchain
Iglc1
Iglv1
G
JHMV 5xFAD vs. JHMV WT – 7dpi
Plin4
Hif3a
Fam107a
Fkbp5
Ighv9−2
Chil3
Gm29650
Cdc25c
Kdm5d
B
E
H
C
JHMV 5xFAD vs. Control 5xFAD Enriched Downregulated Pathways
Cholesterol biosynthesis
Steroid biosynthesis
Neuronal system
Terpenoid backbone biosynthesis
Transmission across chemical synapses
Lipid and lipoprotein metabolism
SREBF and miR-33 in cholesterol and lipid homeostasis
Sodium/chloride-dependent neurotransmitter transporters
Fatty acyl-CoA biosynthesis
Amine compound SLC transporters
JHMV 5xFAD vs Control 5xFAD Enriched Upregulated Pathways
Interleukin-2 signaling pathway
Immune system
Cytoplasmic ribosomal proteins
T cell receptor regulation of apoptosis
Translation
Influenza viral RNA transcription and replication
Immune system signaling by interferons, interleukins, prolactin, & growth hormones
Influenza infection
Lysosome
Leptin influence on immune response
F
JHMV WT vs. Control WT Enriched Downregulated Pathways
Cholesterol biosynthesis
Steroid biosynthesis
Lipid and lipoprotein metabolism
SREBF and miR-33 in cholesterol and lipid homeostasis
SREBP control of lipid biosynthesis
Statin pathway
TGF-beta regulation of extracellular matrix
Terpenoid backbone biosynthesis
Oncostatin M
Sodium/chloride-dependent neurotransmitter transporters
JHMV WT vs. Control WT Enriched Upregulated Pathways
Immune system
Interleukin-2 signaling pathway
Cytoplasmic ribosomal proteins
Immune system signaling by interferons, interleukins, prolactin, and growth hormones
T cell receptor regulation of apoptosis
Translation
Influenza viral RNA transcription and replication
Interferon signaling
Influenza infection
Adaptive immune system
I
JHMV 5xFAD vs JHMV WT Enriched Downregulated Pathways
BDNF signaling pathway
GPCR ligand binding
Gastrin-CREB signaling pathway via PKC and MAPK
Peptide G-protein coupled receptors
G alpha (i) signaling events
Activation of chaperones by IRE1 alpha
Aquaporin-mediated transport
Parkin role in the ubiquitin-proteasomal pathway
Activation of chaperones by ATF6 alpha
Protein processing in the endoplasmic reticulum
JHMV 5xFAD vs JHMV WT Enriched Upregulated Pathways
Hemostasis pathway
Glycosaminoglycan degradation
Oncostatin M
Urokinase-type plasminogen activator (uPA) and uPAR-mediated signaling
Response to elevated platelet cytosolic calcium
Platelet activation, signaling and aggregation
Cell surface interactions at the vascular wall
Hyaluronan metabolism
Interleukin-6 regulation of target genes
HIF-2-alpha transcription factor network
Bulk RNA sequencing: 14 days post-infection
J
JHMV 5xFAD vs. Control 5xFAD – 14dpi
Gpnmb
H2−Q7
Cybb
Psmb8
H2−T23
Lyz2
Gbp2
Igtp
H2−D1
Spp1
Iigp1
B2m
Il12rb1
Ifi47
Ccl5
Ly6c2
Igkc
Igha
Iglc1
M
JHMV WT vs. CTL WT – 14dpi
H2−D1
Ctss
H2−K1
Cd74
Gpnmb
H2−Ab1
Mpeg1
Lyz2
Tap1
H2−Eb1
B2m
C4b
Grn
Laptm5
Trem2
Ctsd
Lgals3bp
Vim
C1qb
C1qa
Cd68
C1qc
H2−Aa
Igtp
H2−Q4
Cybb
H2−T23
Ctsb
Irgm1
Irf7
Ctsz
C3
Igkc
Clec7a
Hmgcs1
Igha
Jchain
Iglc1
Ighg2b
Iglv1
Ighv1−54
P
JHMV 5xFAD vs. JHMV WT – 14dpi
Sumo2
Thy1
Siglec1
Gm10241
K
N
Q
L
JHMV 5xFAD vs. Control 5xFAD Enriched Downregulated Pathways
Cholesterol biosynthesis
Axon guidance
L1CAM interactions
Ion channel transport
Neuroactive ligand-receptor interaction
Steroid biosynthesis
Developmental biology
Interaction between L1-type proteins and ankyrins
Terpenoid backbone biosynthesis
Ligand-gated ion channel transport
JHMV 5xFAD vs Control 5xFAD Enriched Upregulated Pathways
Cytoplasmic ribosomal proteins
Influenza viral RNA transcription and replication
Translation
Influenza infection
Cap-dependent translation initiation
T cell receptor regulation of apoptosis
Immune system
Interleukin-2 signaling pathway
Disease
Activation of mRNA upon binding of the cap-binding complex and eIFs, and subsequent binding to 43S
O
JHMV WT vs. Control WT Enriched Downregulated Pathways
Cholesterol biosynthesis
Neuronal system
Transmission across chemical synapses
Terpenoid backbone biosynthesis
Calcium regulation in the cardiac cell
Steroid biosynthesis
G alpha (i) signaling events
Axon guidance
Myometrial relaxation and contraction pathways
Calcium signaling pathway
JHMV WT vs. Control WT Enriched Upregulated Pathways
Interleukin-2 signaling pathway
Immune system
Cytoplasmic ribosomal proteins
Translation
Influenza viral RNA transcription and replication
T cell receptor regulation of apoptosis
Immune system signaling by interferons, interleukins, prolactin, and growth hormones
Influenza infection
Cap-dependent translation initiation
Lysosome
R
JHMV 5xFAD vs JHMV WT Enriched Downregulated Pathways
Basic mechanisms of SUMOylation
Regular glucocorticoid receptor pathway
Cell adhesion molecules (CAMs)
JHMV 5xFAD vs JHMV WT Enriched Upregulated Pathways
T helper cell surface molecules
Integrin beta-2 pathway
Alpha-M beta-2 integrin signaling
Beta-3 integrin cell surface interactions
Interleukin-4 signaling pathway
Leukocyte transendothelial migration
FSH regulation of apoptosis
NS
p-value
Log2 FC
p-value and Log2 FC
Row minimum
Row maximum
Control WT
Control 5xFAD
JHMV-infected WT
JHMV-infected 5xFAD

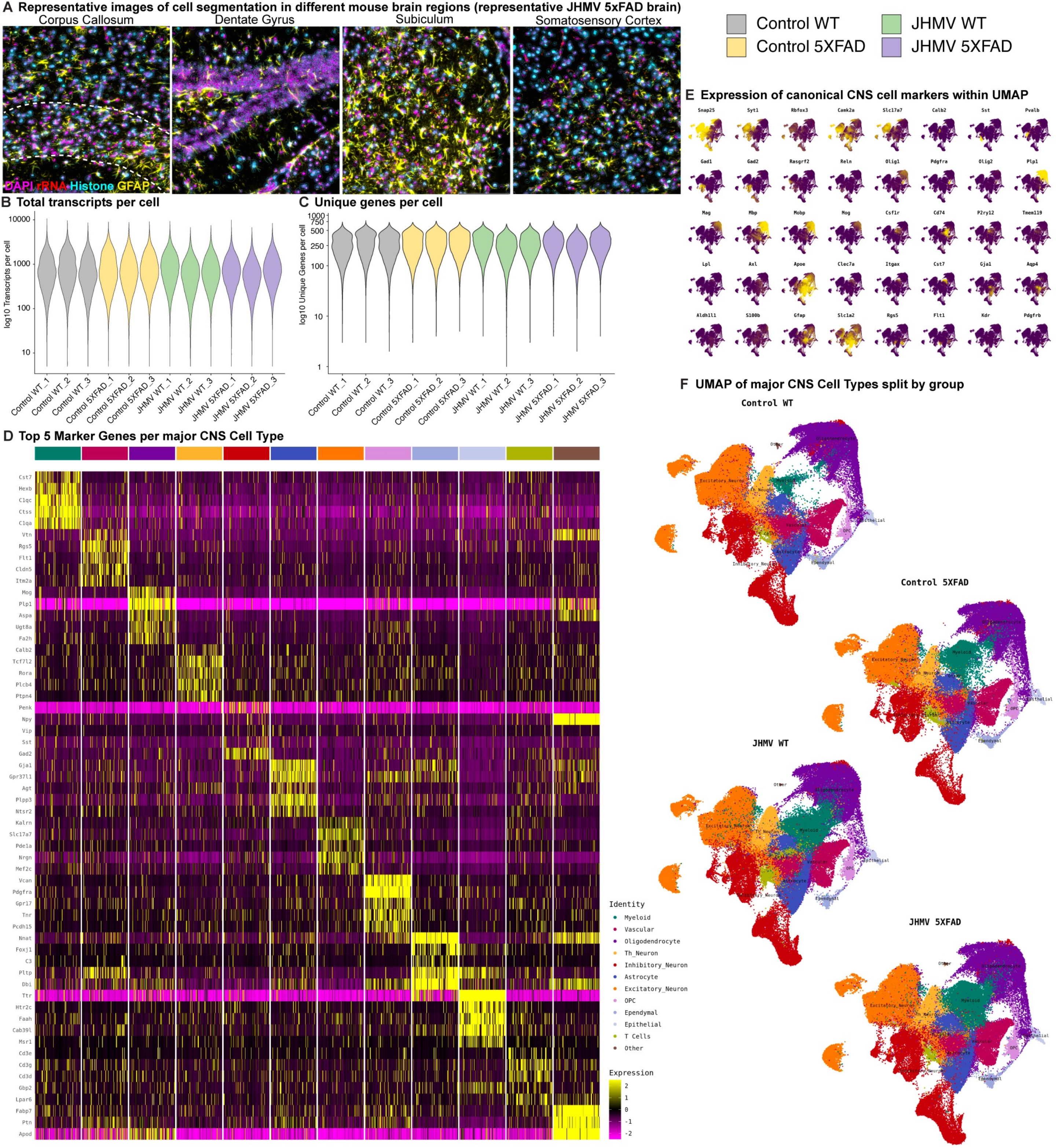
A Representative images of cell segmentation in different mouse brain regions (representative JHMV 5xFAD brain)
Corpus Callosum
Dentate Gyrus
Subiculum
Somatosensory Cortex
DAPI rRNA Histone GFAP
Control WT
JHMV WT
Control 5XFAD
JHMV 5XFAD
B Total transcripts per cell
log10 Transcripts per cell
10000
1000
100
10
C Unique genes per cell
log10 Unique Genes per cell
1000
750
500
250
100
10
1
Control WT_1
Control WT_2
Control WT_3
Control 5XFAD_1
Control 5XFAD_2
Control 5XFAD_3
JHMV WT_1
JHMV WT_2
JHMV WT_3
JHMV 5XFAD_1
JHMV 5XFAD_2
JHMV 5XFAD_3
E Expression of canonical CNS cell markers within UMAP
Snap25
Syt1
Rbfox3
Camk2a
Slc17a7
Calb2
Sst
Pvalb
Gad1
Gad2
Rasgrf2
Reln
Olig1
Pdgfra
Olig2
Plp1
Mag
Mbp
Mobp
Mog
Csf1r
Cd74
P2ry12
Tmem119
Lpl
Axl
Apoe
Clec7a
Itgax
Cst7
Gja1
Aqp4
Aldh1l1
S100b
Gfap
Slc1a2
Rgs5
Flt1
Kdr
Pdgfrb
D Top 5 Marker Genes per major CNS Cell Type
Cst7
Hexb
C1qc
Ctss
C1qa
Vtn
Rgs5
Flt1
Cldn5
Itm2a
Mog
Plp1
Aspa
Ugt8a
Fa2h
Calb2
Tcf7l2
Rora
Plcb4
Ptpn4
Penk
Npy
Vip
Sst
Gad2
Gja1
Gpr37l1
Agt
Plpp3
Ntsr2
Kalrn
Slc17a7
Pde1a
Nrgn
Mef2c
Vcan
Pdgfra
Gpr17
Tnr
Pcdh15
Nnat
Foxj1
C3
Pltp
Dbi
Ttr
Htr2c
Faah
Cab39l
Msr1
Cd3e
Cd3g
Cd3d
Gbp2
Lpar6
Fabp7
Ptn
Apod
Identity
Myeloid
Vascular
Oligodendrocyte
Th_Neuron
Inhibitory_Neuron
Astrocyte
Excitatory_Neuron
OPC
Ependymal
Epithelial
T Cells
Other
Expression
2
1
0
-1
-2
F UMAP of major CNS Cell Types split by group
Control WT
Control 5XFAD
JHMV WT
JHMV 5XFAD

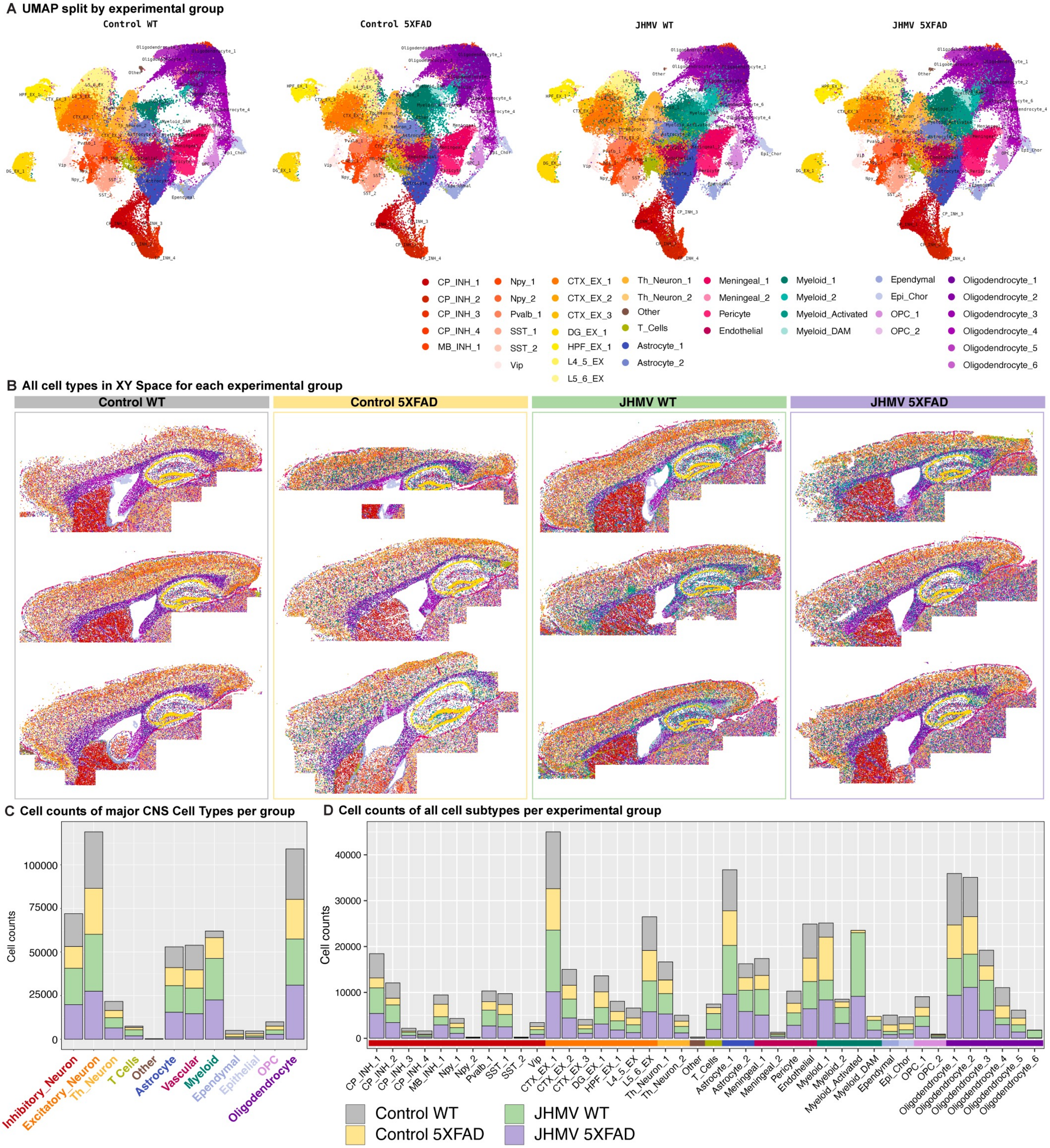

A UMAP split by experimental group
Control WT
Control 5XFAD
JHMV WT
JHMV 5XFAD
CP_INH_1
CP_INH_2
CP_INH_3
CP_INH_4
MB_INH_1
Npy_1
Npy_2
Pvalb_1
SST_1
SST_2
Vip
CTX_EX_1
CTX_EX_2
CTX_EX_3
DG_EX_1
HPF_EX_1
L4_5_EX
L5_6_EX
Th_Neuron_1
Th_Neuron_2
Other
T_Cells
Astrocyte_1
Astrocyte_2
Meningeal_1
Meningeal_2
Pericyte
Endothelial
Myeloid_1
Myeloid_2
Myeloid_Activated
Myeloid_DAM
Ependymal
Epi_Chor
OPC_1
OPC_2
Oligodendrocyte_1
Oligodendrocyte_2
Oligodendrocyte_3
Oligodendrocyte_4
Oligodendrocyte_5
Oligodendrocyte_6
B All cell types in XY Space for each experimental group
Control WT
Control 5XFAD
JHMV WT
JHMV 5XFAD
C Cell counts of major CNS Cell Types per group
Cell counts
100000
75000
50000
25000
0
Inhibitory_Neuron
Excitatory_Neuron
Th_Neuron
T Cells
Other
Astrocyte
Vascular
Myeloid
Ependymal
Epithelial
OPC
Oligodendrocyte
D Cell counts of all cell subtypes per experimental group
Cell counts
40000
30000
20000
10000
0
CP_INH_1
CP_INH_2
CP_INH_3
CP_INH_4
MB_INH_1
Npy_1
Npy_2
Pvalb_1
SST_1
SST_2
Vip
CTX_EX_1
CTX_EX_2
CTX_EX_3
DG_EX_1
HPF_EX_1
L4_5_EX
L5_6_EX
Th_Neuron_1
Th_Neuron_2
Other
T_Cells
Astrocyte_1
Astrocyte_2
Meningeal_1
Meningeal_2
Pericyte
Endothelial
Myeloid_1
Myeloid_2
Myeloid_Activated
Myeloid_DAM
Ependymal
Epi_Chor
OPC_1
OPC_2
Oligodendrocyte_1
Oligodendrocyte_2
Oligodendrocyte_3
Oligodendrocyte_4
Oligodendrocyte_5
Oligodendrocyte_6
Control WT
Control 5XFAD
JHMV WT
JHMV 5XFAD

#### DGE Analysis for JHMV WT vs. Control WT: All CNS Subtypes

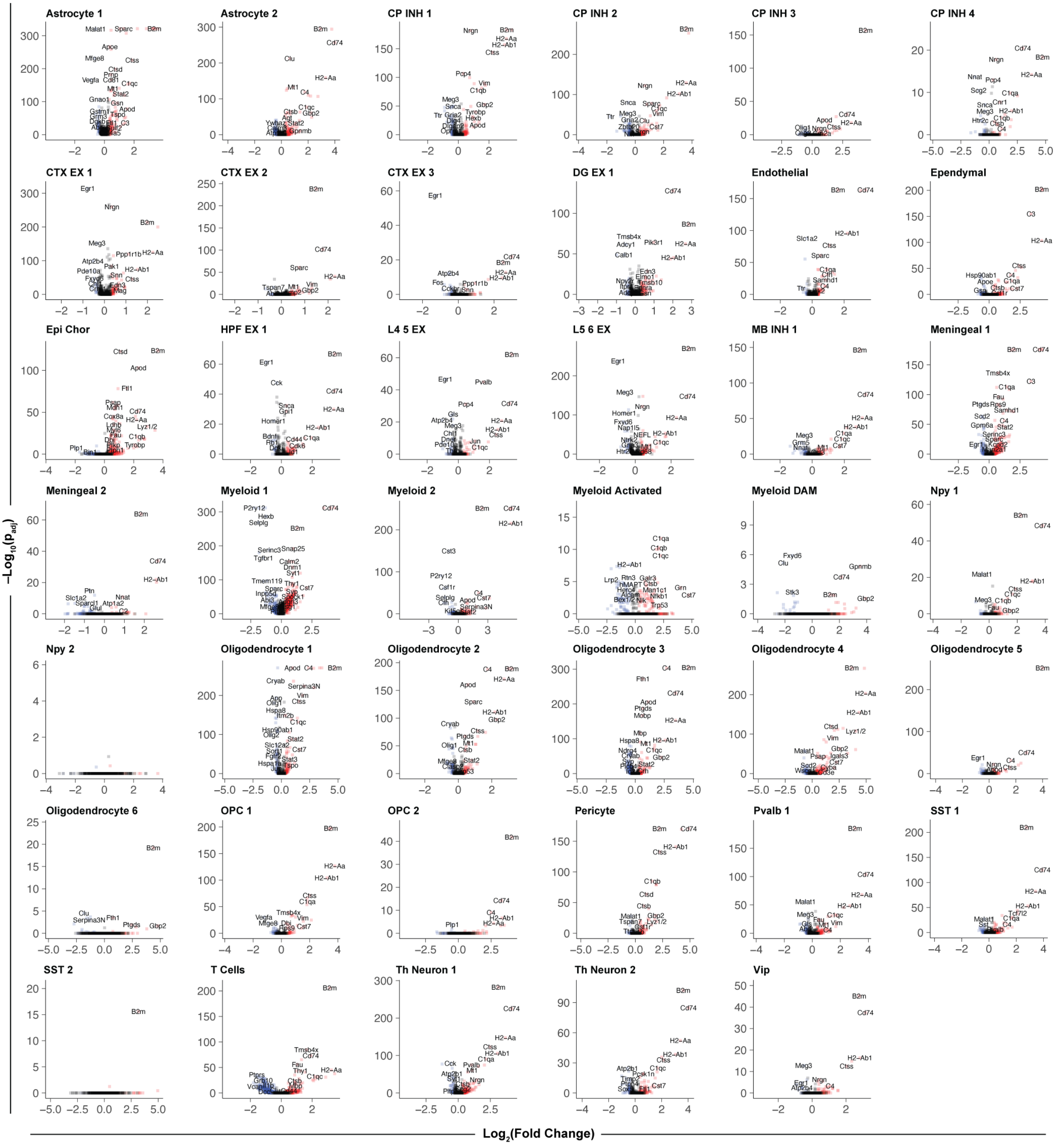

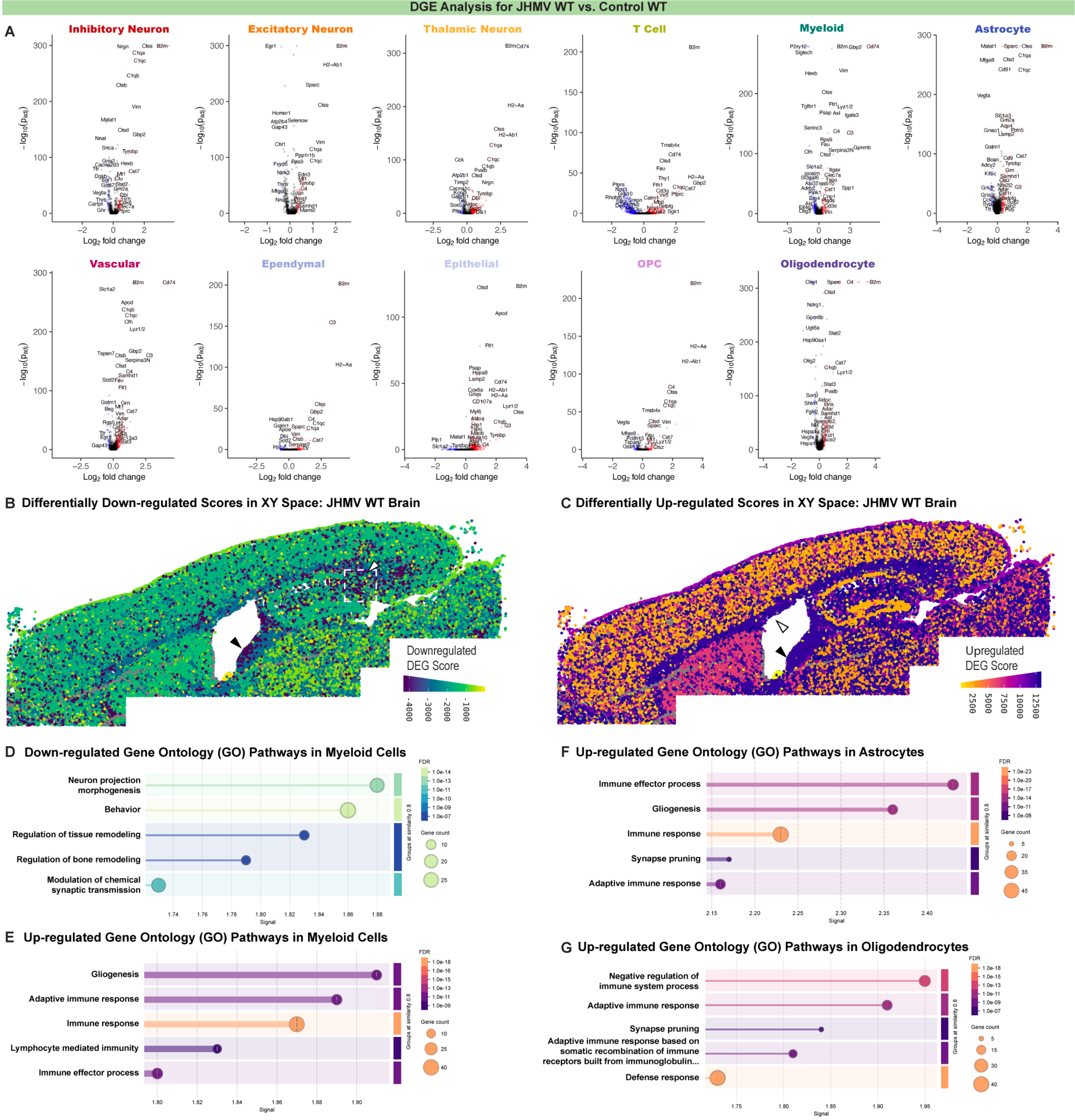
DGE Analysis for JHMV WT vs. Control WT
A
Inhibitory Neuron
Excitatory Neuron
Thalamic Neuron
T Cell
Myeloid
Astrocyte
Vascular
Ependymal
Epithelial
OPC
Oligodendrocyte
−log10(Padj)
Log2 fold change
B Differentially Down-regulated Scores in XY Space: JHMV WT Brain
Downregulated DEG Score
-4000
-3000
-2000
-1000
C Differentially Up-regulated Scores in XY Space: JHMV WT Brain
Upregulated DEG Score
2500
5000
7500
10000
12500
D Down-regulated Gene Ontology (GO) Pathways in Myeloid Cells
Neuron projection morphogenesis
Behavior
Regulation of tissue remodeling
Regulation of bone remodeling
Modulation of chemical synaptic transmission
Signal
FDR
Gene count
Groups at similarity 0.8
E Up-regulated Gene Ontology (GO) Pathways in Myeloid Cells
Gliogenesis
Adaptive immune response
Immune response
Lymphocyte mediated immunity
Immune effector process
Signal
F Up-regulated Gene Ontology (GO) Pathways in Astrocytes
Immune effector process
Gliogenesis
Immune response
Synapse pruning
Adaptive immune response
Signal
G Up-regulated Gene Ontology (GO) Pathways in Oligodendrocytes
Negative regulation of immune system process
Adaptive immune response
Synapse pruning
Adaptive immune response based on somatic recombination of immune receptors built from immunoglobulin...
Defense response
Signal

#### DGE Analysis for Control 5XFAD vs. Control WT: All CNS Subtypes

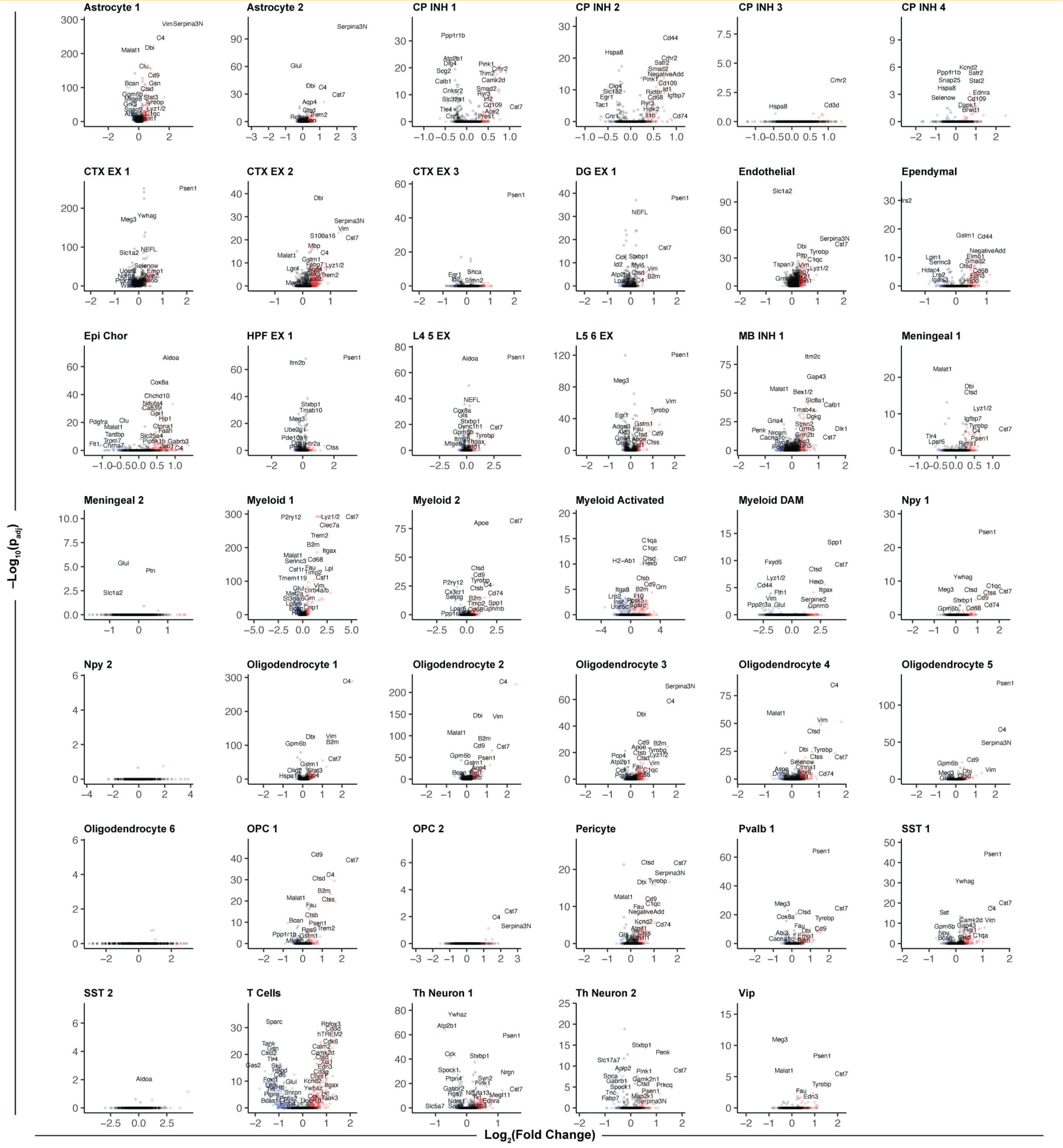

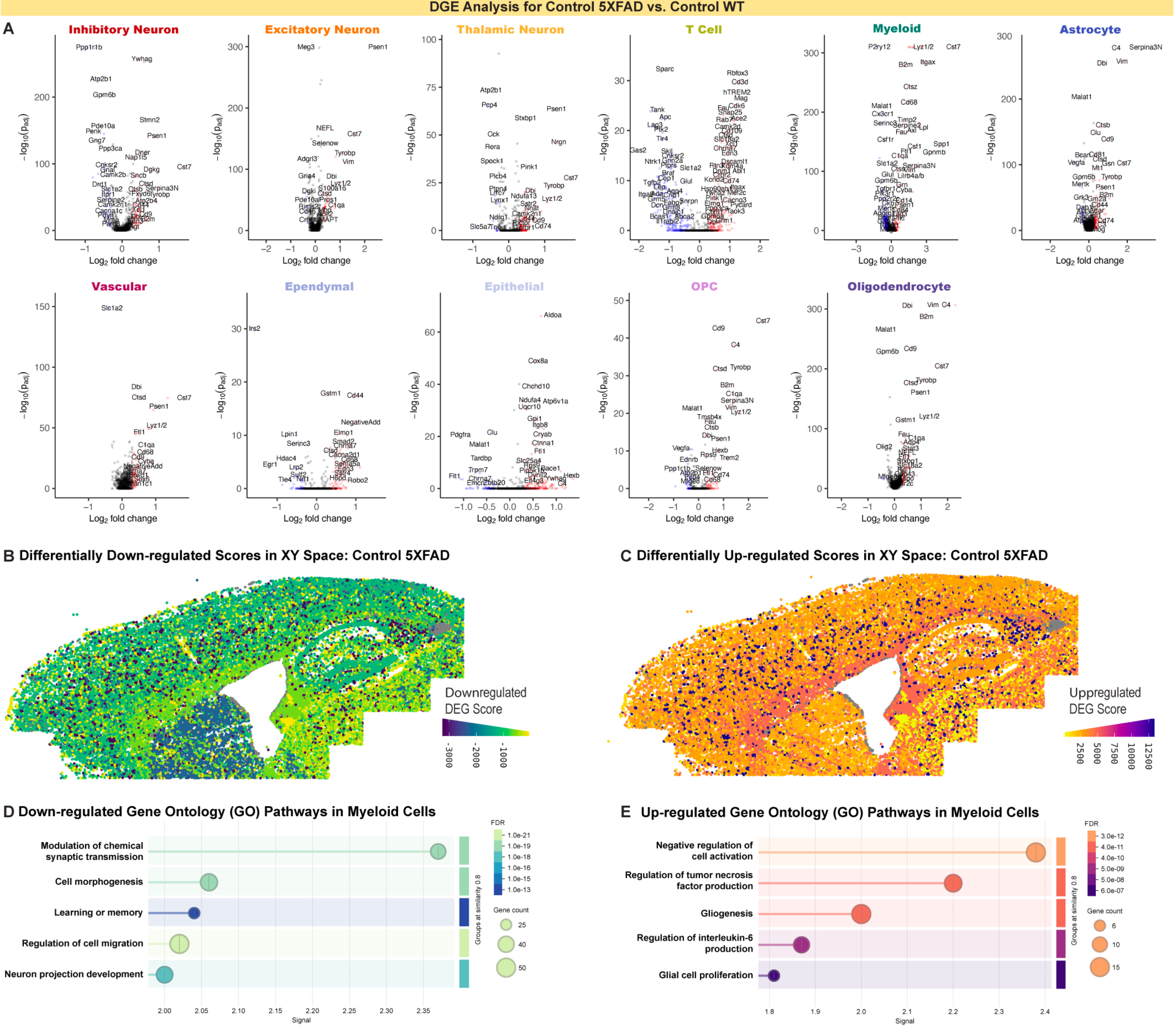
DGE Analysis for Control 5XFAD vs. Control WT
A
Inhibitory Neuron
Excitatory Neuron
Thalamic Neuron
T Cell
Myeloid
Astrocyte
Vascular
Ependymal
Epithelial
OPC
Oligodendrocyte
Log2 fold change
−log10(padj)
B Differentially Down-regulated Scores in XY Space: Control 5XFAD
Downregulated DEG Score
-3000
-2000
-1000
C Differentially Up-regulated Scores in XY Space: Control 5XFAD
Upregulated DEG Score
2500
5000
7500
10000
12500
D Down-regulated Gene Ontology (GO) Pathways in Myeloid Cells
Modulation of chemical synaptic transmission
Cell morphogenesis
Learning or memory
Regulation of cell migration
Neuron projection development
Signal
FDR
Gene count
Groups at similarity 0.8
E Up-regulated Gene Ontology (GO) Pathways in Myeloid Cells
Negative regulation of cell activation
Regulation of tumor necrosis factor production
Gliogenesis
Regulation of interleukin-6 production
Glial cell proliferation
Signal
FDR
Gene count
Groups at similarity 0.8

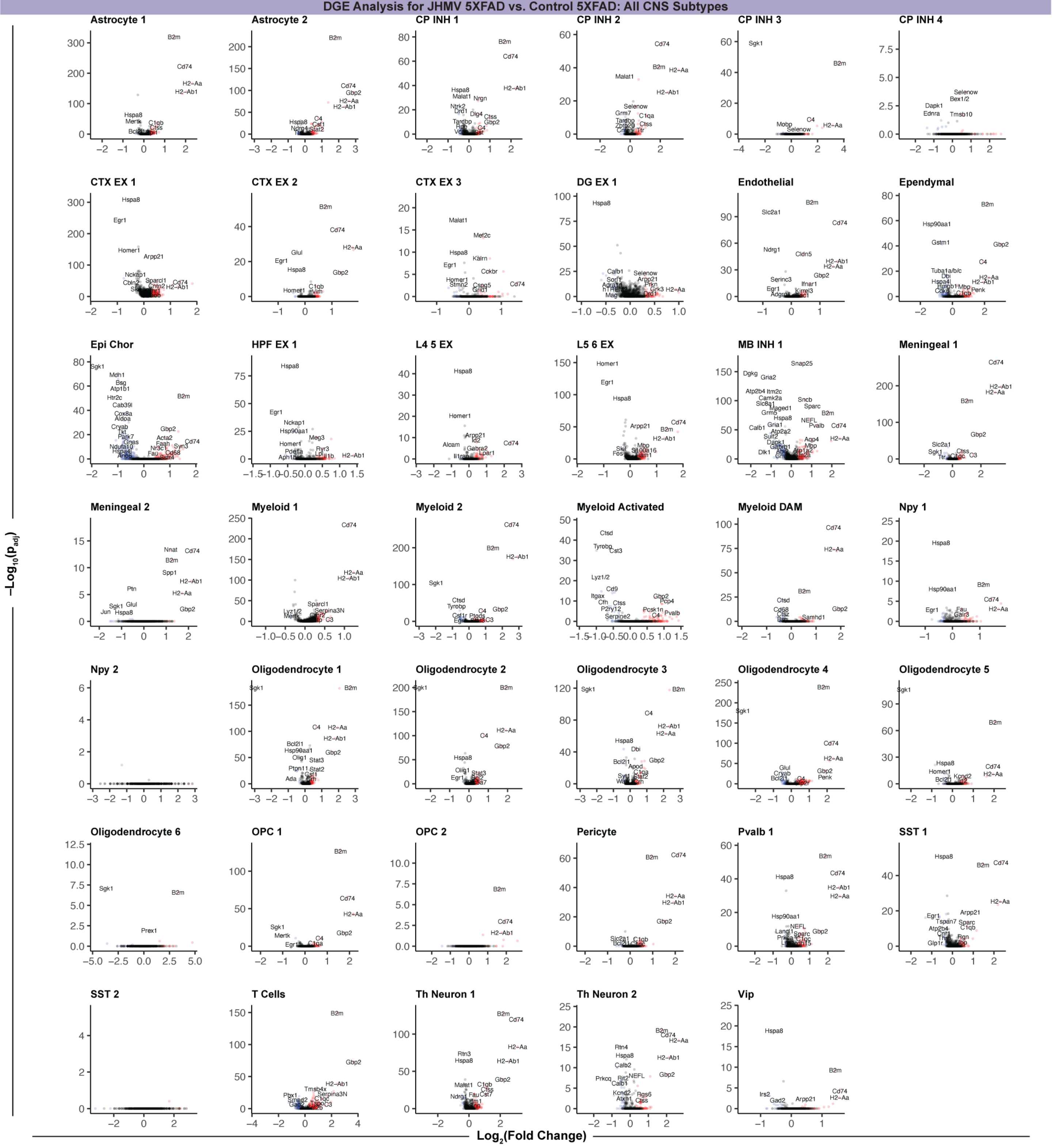
DGE Analysis for JHMV 5XFAD vs. Control 5XFAD: All CNS Subtypes
Astrocyte 1
Astrocyte 2
CP INH 1
CP INH 2
CP INH 3
CP INH 4
CTX EX 1
CTX EX 2
CTX EX 3
DG EX 1
Endothelial
Ependymal
Epi Chor
HPF EX 1
L4 5 EX
L5 6 EX
MB INH 1
Meningeal 1
Meningeal 2
Myeloid 1
Myeloid 2
Myeloid Activated
Myeloid DAM
Npy 1
Npy 2
Oligodendrocyte 1
Oligodendrocyte 2
Oligodendrocyte 3
Oligodendrocyte 4
Oligodendrocyte 5
Oligodendrocyte 6
OPC 1
OPC 2
Pericyte
Pvalb 1
SST 1
SST 2
T Cells
Th Neuron 1
Th Neuron 2
Vip
$-Log_{10}(p_{adj})$
$Log_2$(Fold Change)

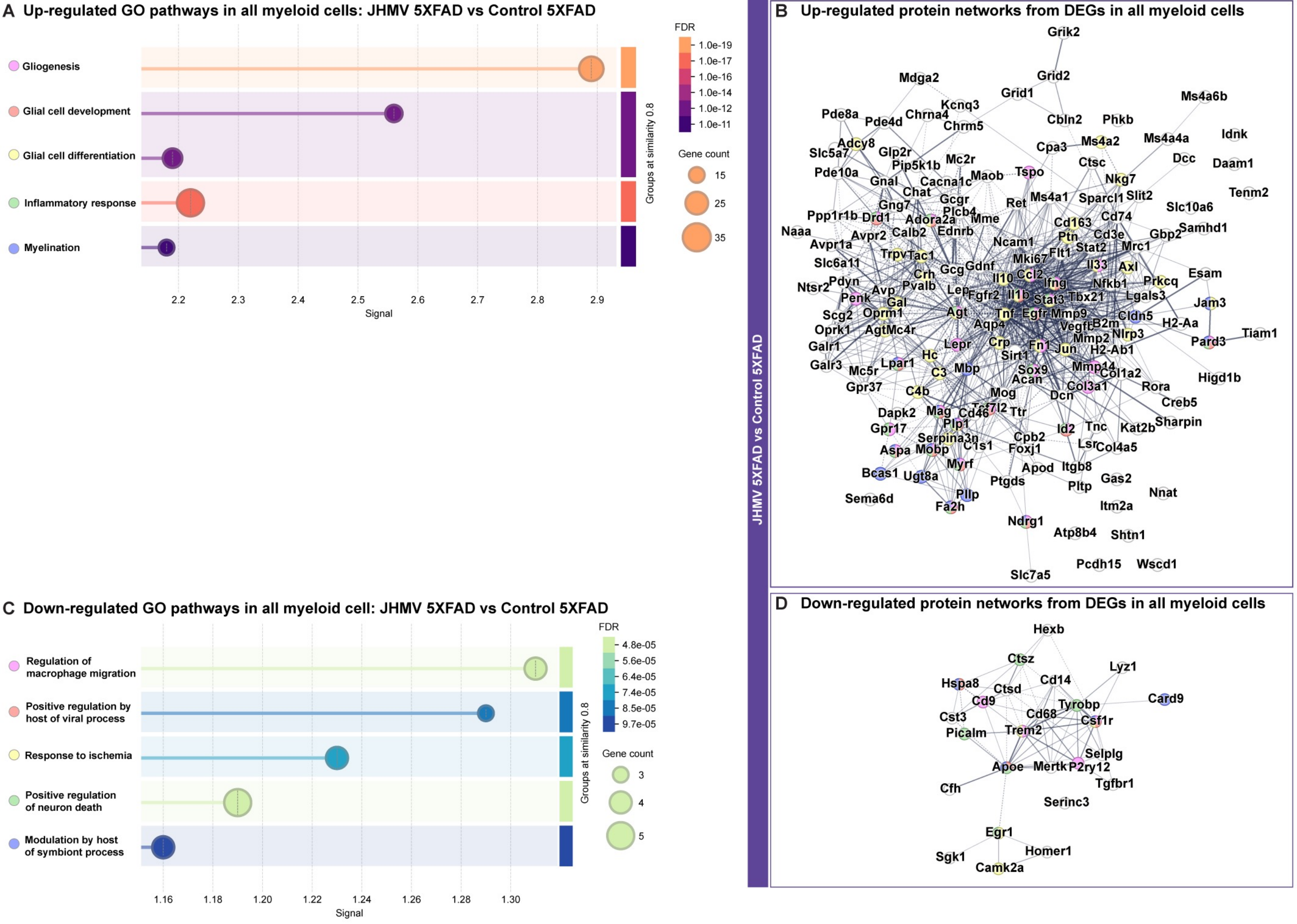

A Up-regulated GO pathways in all myeloid cells: JHMV 5XFAD vs Control 5XFAD
Gliogenesis
Glial cell development
Glial cell differentiation
Inflammatory response
Myelination
Signal
2.2
2.3
2.4
2.5
2.6
2.7
2.8
2.9
Groups at similarity 0.8
FDR
1.0e-19
1.0e-17
1.0e-16
1.0e-14
1.0e-12
1.0e-11
Gene count
15
25
35
B Up-regulated protein networks from DEGs in all myeloid cells
JHMV 5XFAD vs Control 5XFAD
Grik2
Mdga2
Grid2
Grid1
Kcnq3
Pde8a
Pde4d
Chrna4
Chrm5
Cbln2
Phkb
Ms4a6b
Adcy8
Slc5a7
Glp2r
Cpa3
Ms4a2
Ms4a4a
Idnk
Pip5k1b
Mc2r
Ctsc
Dcc
Daam1
Pde10a
Tspo
Gnal
Maob
Nkg7
Cacna1c
Chat
Ms4a1
Sparcl1
Slit2
Tenm2
Gng7
Gcgr
Ret
Slc10a6
Ppp1r1b
Drd1
Adora2a
Plcb4
Mme
Cd163
Cd74
Samhd1
Naaa
Calb2
Ednrb
Gbp2
Avpr2
Ptn
Cd3e
Avpr1a
Ncam1
Stat2
Mrc1
Trpv1
Tac1
Flt1
Slc6a11
Mki67
Il33
Axl
Gcg
Gdnf
Crh
Ccl2
Esam
Il10
Ifng
Prkcq
Ntsr2
Pdyn
Pvalb
Nfkb1
Avp
Lep
Fgfr2
Penk
Il1b
Stat3
Tbx21
Lgals3
Gal
Jam3
Scg2
Oprm1
Agt
Tnf
Egfr
Mmp9
Cldn5
Aqp4
Vegfb
B2m
H2-Aa
Oprk1
Agt
Mc4r
Nlrp3
Tiam1
Mmp2
Pard3
Galr1
Lepr
Crp
Fn1
Jun
H2-Ab1
Hc
Sirt1
Galr3
Lpar1
Mc5r
C3
Mbp
Sox9
Mmp14
Col1a2
Higd1b
Acan
Gpr37
Col3a1
Rora
C4b
Mog
Dcn
Creb5
Tcf7l2
Dapk2
Mag
Cd46
Ttr
Sharpin
Plp1
Gpr17
Tnc
Kat2b
Id2
Serpina3n
Cpb2
Aspa
Mobp
C1s1
Foxj1
Lsr
Col4a5
Myrf
Itgb8
Bcas1
Ugt8a
Apod
Gas2
Ptgds
Pltp
Pllp
Nnat
Sema6d
Itm2a
Fa2h
Ndrg1
Atp8b4
Shtn1
Pcdh15
Wscd1
Slc7a5
C Down-regulated GO pathways in all myeloid cell: JHMV 5XFAD vs Control 5XFAD
Regulation of macrophage migration
Positive regulation by host of viral process
Response to ischemia
Positive regulation of neuron death
Modulation by host of symbiont process
Signal
1.16
1.18
1.20
1.22
1.24
1.26
1.28
1.30
Groups at similarity 0.8
FDR
4.8e-05
5.6e-05
6.4e-05
7.4e-05
8.5e-05
9.7e-05
Gene count
3
4
5
D Down-regulated protein networks from DEGs in all myeloid cells
Hexb
Ctsz
Lyz1
Hspa8
Cd14
Ctsd
Cd9
Card9
Tyrobp
Cst3
Cd68
Csf1r
Picalm
Trem2
Selplg
Apoe
Mertk
P2ry12
Cfh
Tgfbr1
Serinc3
Egr1
Homer1
Sgk1
Camk2a

### Myeloid clusters: Spatial transcriptomics analysis, Top 10 Expressed Genes, and Location in XY space

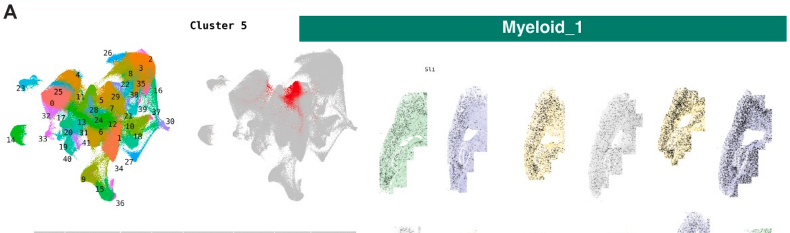


| | p_val | avg_log2FC | pct.1 | pct.2 | p_val_adj | cluster | gene | pct_diff |
|---|---|---|---|---|---|---|---|---|
| 4373 | 0 | 3.225924 | 0.828 | 0.191 | 0 | 5 | Hexb | 0.637 |
| 4374 | 0 | 3.204947 | 0.689 | 0.166 | 0 | 5 | Tyrobp | 0.523 |
| 4375 | 0 | 3.199240 | 0.469 | 0.103 | 0 | 5 | Cst7 | 0.366 |
| 4376 | 0 | 2.990977 | 0.438 | 0.084 | 0 | 5 | Trem2 | 0.354 |
| 4377 | 0 | 2.947652 | 0.700 | 0.161 | 0 | 5 | Csf1r | 0.539 |
| 4378 | 0 | 2.938707 | 0.413 | 0.076 | 0 | 5 | Selplg | 0.337 |
| 4379 | 0 | 2.773960 | 0.531 | 0.125 | 0 | 5 | Cx3cr1 | 0.406 |
| 4380 | 0 | 2.769211 | 0.843 | 0.206 | 0 | 5 | C1qc | 0.637 |
| 4381 | 0 | 2.645798 | 0.867 | 0.242 | 0 | 5 | C1qa | 0.625 |
| 4382 | 0 | 2.633358 | 0.969 | 0.674 | 0 | 5 | Ctsd | 0.295 |

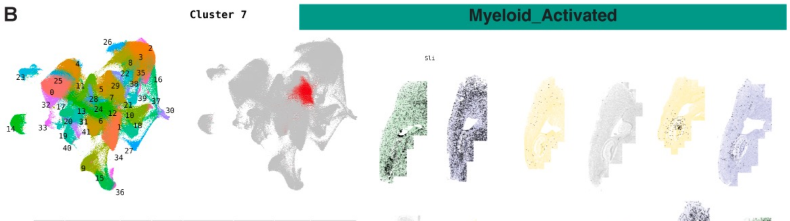


| | p_val | avg_log2FC | pct.1 | pct.2 | p_val_adj | cluster | gene | pct_diff |
|---|---|---|---|---|---|---|---|---|
| 6092 | 0 | 4.269159 | 0.840 | 0.125 | 0 | 7 | H2-Ab1 | 0.715 |
| 6093 | 0 | 4.227990 | 0.848 | 0.123 | 0 | 7 | H2-Aa | 0.725 |
| 6094 | 0 | 4.169679 | 0.992 | 0.272 | 0 | 7 | Cd74 | 0.720 |
| 6095 | 0 | 3.162702 | 0.939 | 0.284 | 0 | 7 | Ctss | 0.655 |
| 6096 | 0 | 3.107893 | 0.872 | 0.244 | 0 | 7 | C1qa | 0.628 |
| 6097 | 0 | 3.067096 | 0.842 | 0.208 | 0 | 7 | C1qc | 0.634 |
| 6098 | 0 | 2.874776 | 0.888 | 0.314 | 0 | 7 | C1qb | 0.574 |
| 6099 | 0 | 2.777657 | 0.753 | 0.197 | 0 | 7 | Hexb | 0.556 |
| 6100 | 0 | 2.603173 | 0.645 | 0.165 | 0 | 7 | Csf1r | 0.480 |
| 6101 | 0 | 2.512817 | 0.370 | 0.079 | 0 | 7 | Selplg | 0.291 |

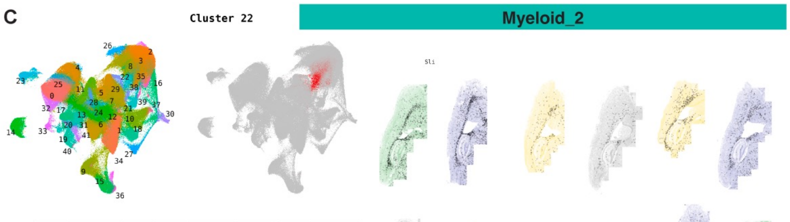


| | p_val | avg_log2FC | pct.1 | pct.2 | p_val_adj | cluster | gene | pct_diff |
|---|---|---|---|---|---|---|---|---|
| 18088 | 0 | 2.150403 | 0.948 | 0.303 | 0 | 22 | Ctss | 0.645 |
| 18089 | 0 | 1.990397 | 0.825 | 0.227 | 0 | 22 | C1qc | 0.598 |
| 18090 | 0 | 1.989139 | 0.758 | 0.297 | 0 | 22 | Cd74 | 0.461 |
| 18091 | 0 | 1.983204 | 0.869 | 0.263 | 0 | 22 | C1qa | 0.606 |
| 18092 | 0 | 1.892670 | 0.554 | 0.149 | 0 | 22 | H2-Aa | 0.405 |
| 18093 | 0 | 1.833196 | 0.678 | 0.184 | 0 | 22 | Mog | 0.494 |
| 18094 | 0 | 1.829746 | 0.880 | 0.331 | 0 | 22 | C1qb | 0.549 |
| 18095 | 0 | 1.825003 | 0.999 | 0.568 | 0 | 22 | Plp1 | 0.431 |
| 18096 | 0 | 1.816731 | 0.711 | 0.214 | 0 | 22 | Hexb | 0.497 |
| 18097 | 0 | 1.786108 | 0.530 | 0.151 | 0 | 22 | H2-Ab1 | 0.379 |

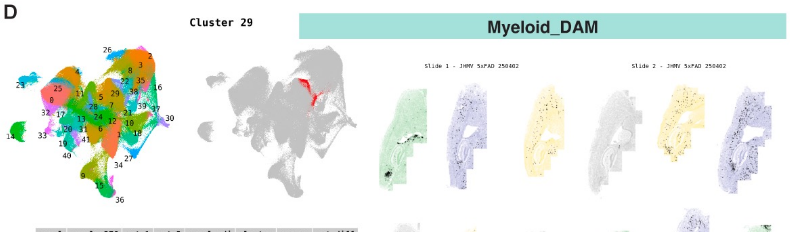


| | p_val | avg_log2FC | pct.1 | pct.2 | p_val_adj | cluster | gene | pct_diff |
|---|---|---|---|---|---|---|---|---|
| 23705 | 0 | 6.442018 | 0.751 | 0.063 | 0 | 29 | Spp1 | 0.688 |
| 23706 | 0 | 5.914087 | 0.709 | 0.043 | 0 | 29 | Gpnmb | 0.666 |
| 23707 | 0 | 4.761462 | 0.869 | 0.111 | 0 | 29 | Lyz1/2 | 0.758 |
| 23708 | 0 | 4.311550 | 0.504 | 0.058 | 0 | 29 | lgals3 | 0.446 |
| 23709 | 0 | 4.154825 | 0.673 | 0.116 | 0 | 29 | Cst7 | 0.557 |
| 23710 | 0 | 3.727570 | 0.519 | 0.086 | 0 | 29 | Itgax | 0.433 |
| 23711 | 0 | 3.568844 | 0.889 | 0.185 | 0 | 29 | Tyrobp | 0.704 |
| 23712 | 0 | 3.430195 | 0.612 | 0.097 | 0 | 29 | Trem2 | 0.515 |
| 23713 | 0 | 3.380710 | 0.996 | 0.686 | 0 | 29 | Ctsd | 0.310 |
| 23714 | 0 | 3.268361 | 0.725 | 0.150 | 0 | 29 | H2-Aa | 0.575 |

Control WT | JHMV WT | Control 5XFAD | JHMV 5XFAD

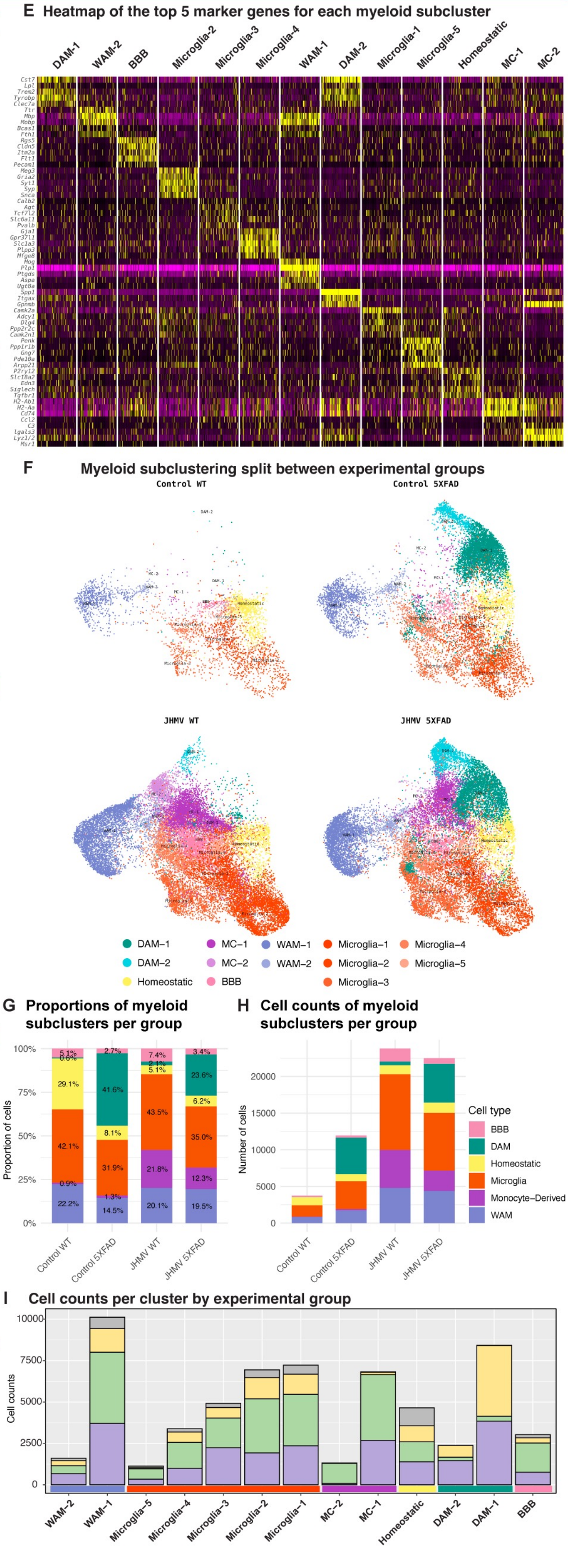

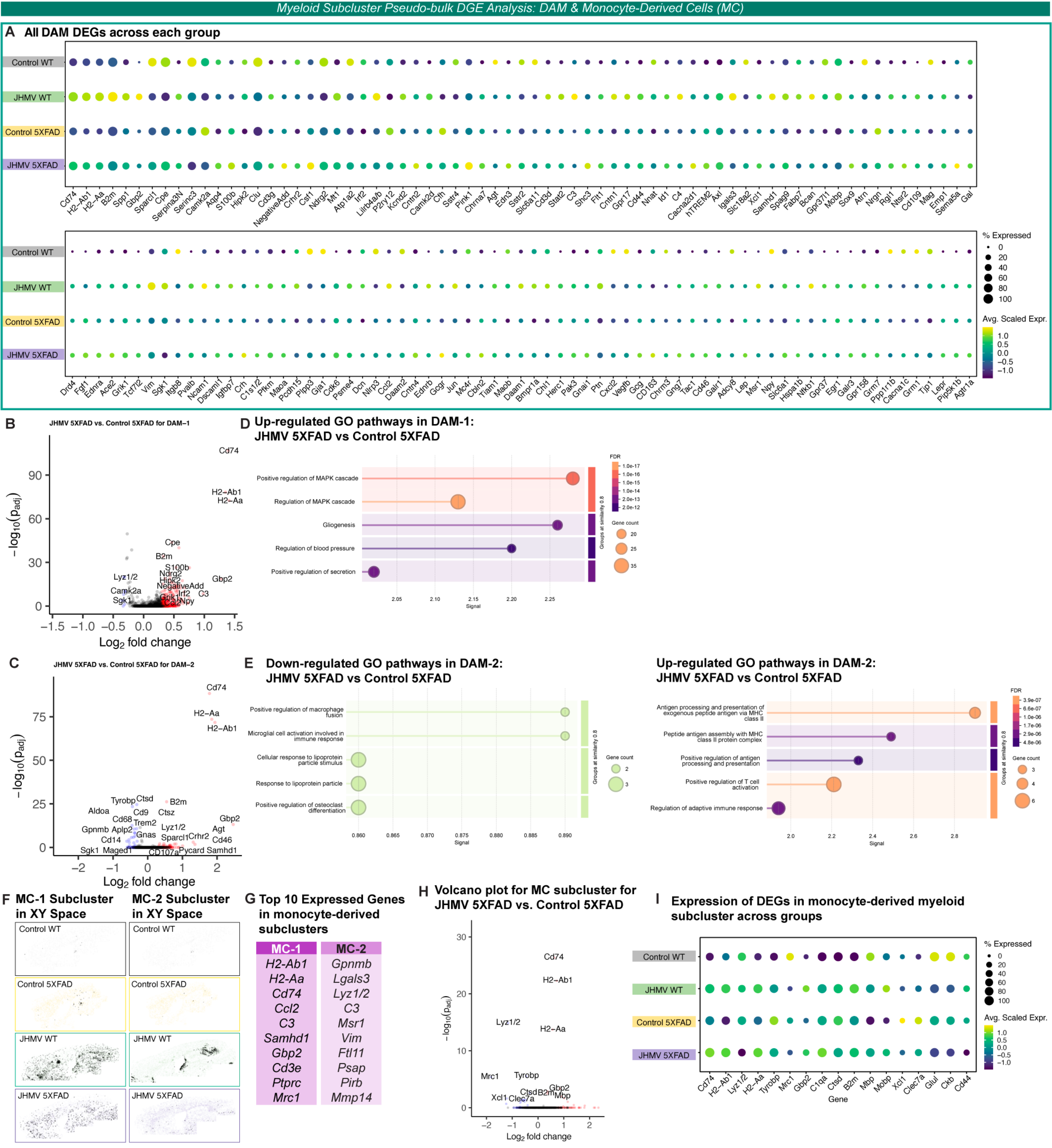
Myeloid Subcluster Pseudo-bulk DGE Analysis: DAM & Monocyte-Derived Cells (MC)
A All DAM DEGs across each group
Control WT
JHMV WT
Control 5XFAD
JHMV 5XFAD
Cd74 H2-Ab1 H2-Aa B2m Spp1 Gbp2 Sparcl1 Cpe Serpina3N Serinc3 Camk2a Aqp4 S100b Hipk2 Clu Cd9 NegativeAdd Crhr2 Csf1 Ndrg2 Mt1 Atp1a2 Irf2 Lilrb4a/b P2ry12 Kcnd2 Cntn2 Camk2d Cfh Sstr4 Pink1 Chrna7 Agt Edn3 Sstr2 Slc6a11 Cd3d Stat2 C3 Shc3 Flt1 Cntn1 Gpr17 Cd44 Nnat Id1 C4 Cacna2d1 hTREM2 Axl Igals3 Slc18a2 Xcl1 Samhd1 Spag9 Fabp7 Bcan Gpr37l1 Mobp Sox9 Atrn Nrgn Rgl1 Ntsr2 Cd109 Mag Emp1 Sema5a Gal
Drd4 Fgf1 Ednra Ace2 Grik1 Tcf7l2 Vim Sgk1 Itgb8 Pvalb Ncam1 Dscaml1 Igfbp7 Crh C1s1/2 Pfkm Maoa Pcdh15 Plpp3 Gja1 Cdk6 Psme4 Dcn Nlrp3 Ccl2 Daam2 Cntn4 Ednrb Gcgr Jun Mc4r Cbln2 Tiam1 Maob Daam1 Bmpr1a Chi1 Herc1 Pak3 Gnai1 Ptn Cxcl2 Vegfb Gcg CD163 Chrm3 Gng7 Tac1 Cd46 Galr1 Adcy8 Lep Msr1 Npy Slc6a1 Hspa1b Nfkb1 Gpr37 Egr1 Galr3 Gpr158 Grm7 Ppp1r1b Cacna1c Grm1 Tlp1 Lepr Pip5k1b Agtr1a
% Expressed
0
20
40
60
80
100
Avg. Scaled Expr.
1.0
0.5
0.0
−0.5
−1.0
B JHMV 5XFAD vs. Control 5XFAD for DAM−1
$-\log_{10}(p_{adj})$
$\text{Log}_2$ fold change
Cd74
H2−Ab1
H2−Aa
Cpe
B2m
S100b
Ndrg2
Hipk2
NegativeAdd
Gbp2
Lyz1/2
Camk2a
Sgk1
Grik1
Irf2
C3
Npy
D Up-regulated GO pathways in DAM-1: JHMV 5XFAD vs Control 5XFAD
Positive regulation of MAPK cascade
Regulation of MAPK cascade
Gliogenesis
Regulation of blood pressure
Positive regulation of secretion
Signal
Groups at similarity 0.8
FDR
Gene count
C JHMV 5XFAD vs. Control 5XFAD for DAM−2
Cd74
H2−Aa
H2−Ab1
Tyrobp
Ctsd
B2m
Aldoa
Cd9
Ctsz
Cd68
Trem2
Gbp2
Gpnmb
Aplp2
Lyz1/2
Agt
Gnas
Sparcl1
Crhr2
Cd14
Cd46
Sgk1
Maged1
CD107a
Pycard
Samhd1
E Down-regulated GO pathways in DAM-2: JHMV 5XFAD vs Control 5XFAD
Positive regulation of macrophage fusion
Microglial cell activation involved in immune response
Cellular response to lipoprotein particle stimulus
Response to lipoprotein particle
Positive regulation of osteoclast differentiation
Up-regulated GO pathways in DAM-2: JHMV 5XFAD vs Control 5XFAD
Antigen processing and presentation of exogenous peptide antigen via MHC class II
Peptide antigen assembly with MHC class II protein complex
Positive regulation of antigen processing and presentation
Positive regulation of T cell activation
Regulation of adaptive immune response
F MC-1 Subcluster in XY Space
MC-2 Subcluster in XY Space
G Top 10 Expressed Genes in monocyte-derived subclusters
MC-1
H2-Ab1
H2-Aa
Cd74
Ccl2
C3
Samhd1
Gbp2
Cd3e
Ptprc
Mrc1
MC-2
Gpnmb
Lgals3
Lyz1/2
C3
Msr1
Vim
Ftl11
Psap
Pirb
Mmp14
H Volcano plot for MC subcluster for JHMV 5XFAD vs. Control 5XFAD
Cd74
H2−Ab1
Lyz1/2
H2−Aa
Mrc1
Tyrobp
Gbp2
Ctsd
B2m
Mbp
Xcl1
Clec7a
I Expression of DEGs in monocyte-derived myeloid subcluster across groups
Cd74 H2-Ab1 Lyz1/2 H2-Aa Tyrobp Mrc1 Gbp2 C1qa Ctsd B2m Mbp Mobp Xcl1 Clec7a Glul Ckb Cd44
Gene

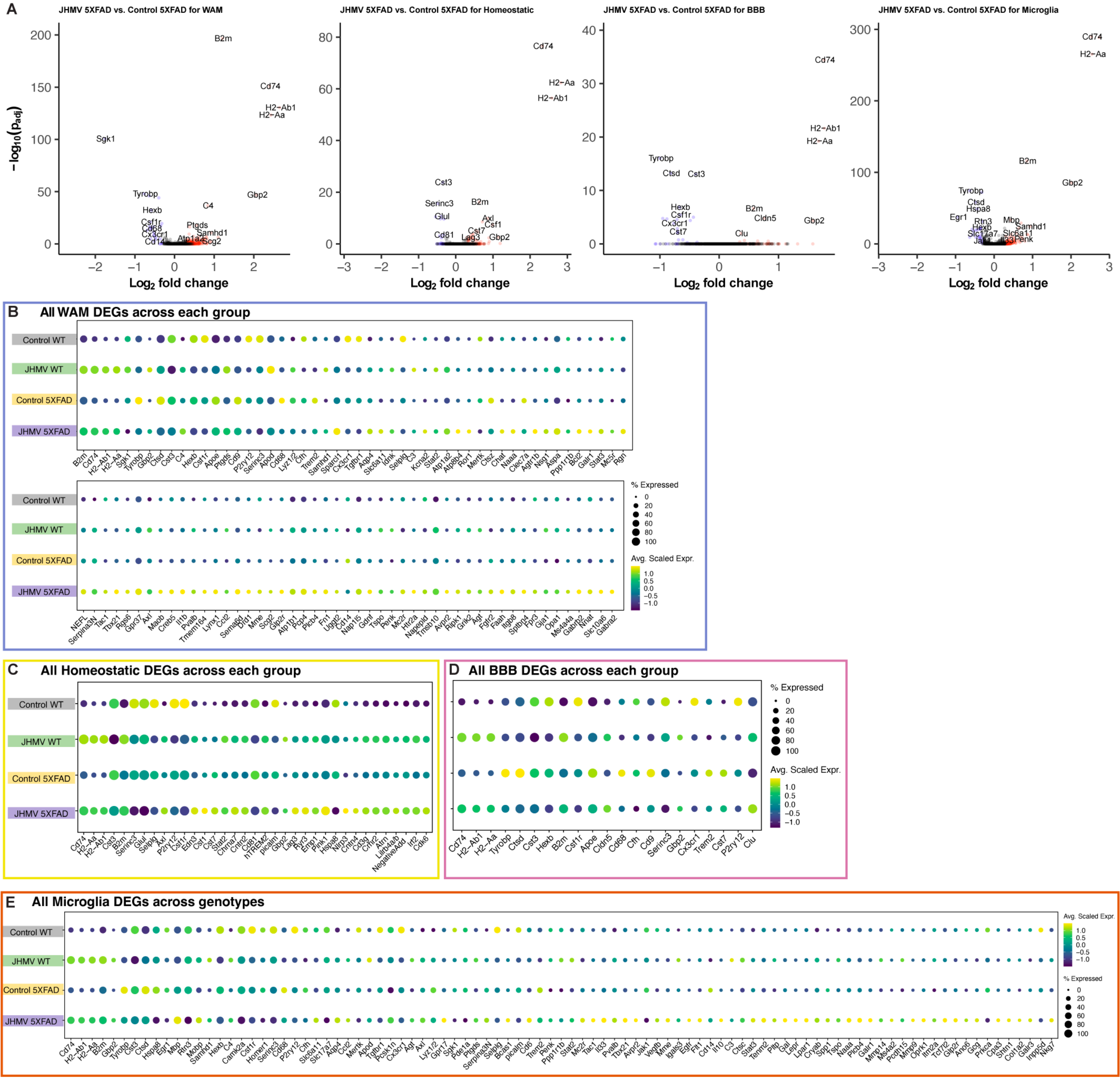

A
JHMV 5XFAD vs. Control 5XFAD for WAM
JHMV 5XFAD vs. Control 5XFAD for Homeostatic
JHMV 5XFAD vs. Control 5XFAD for BBB
JHMV 5XFAD vs. Control 5XFAD for Microglia
−log10(padj)
Log2 fold change
B2m
Cd74
H2−Ab1
H2−Aa
Sgk1
Tyrobp
Hexb
Csf1r
Cd68
Cx3cr1
Cd14
C4
Ptgds
Samhd1
Atp1a2
Scg2
Gbp2
Cst3
Serinc3
Glul
Cd81
Lag3
Cst7
Axl
Csf1
Ctsd
Cldn5
Clu
Hspa8
Egr1
Rtn3
Mbp
Slc17a7
Slc6a11
Penk
B
All WAM DEGs across each group
Control WT
JHMV WT
Control 5XFAD
JHMV 5XFAD
% Expressed
0
20
40
60
80
100
Avg. Scaled Expr.
1.0
0.5
0.0
−0.5
−1.0
C
All Homeostatic DEGs across each group
D
All BBB DEGs across each group
E
All Microglia DEGs across genotypes
